# Dynamic interactions between sensory and decision-making circuits shape temporal evidence weighting

**DOI:** 10.64898/2026.10.09.757813

**Authors:** Citlalli Vivar, Demetrio Ferro, Aaron J. Levi, Alexander C. Huk, Klaus Wimmer

## Abstract

Perceptual decisions require the brain to integrate sensory evidence over time, yet the temporal weighting of evidence can be flexibly adjusted according to task demands. How such flexibility emerges from interactions between sensory and decision circuits remains unclear. Here, we combined behavioral and neural measurements with recurrent network modeling to investigate the mechanisms underlying temporal weighting of sensory evidence. Macaques performed a motion-discrimination task in which the temporal statistics of sensory evidence favored early, late or uniform integration. Behavioral psychophysical kernels revealed corresponding shifts in the influence of evidence across time, accompanied by systematic changes in motion discriminability and choice-related activity modulation in visual area MT. We developed a recurrent sensory-decision network incorporating reciprocal feedforward and feedback interactions and a temporally structured modulatory input, resembling an urgency signal. The model reproduced the observed behavioral and neural signatures across temporal weighting conditions. Mechanistically, shifting the onset and dynamics of modulatory inputs systematically reconfigured the temporal profile of evidence weighting without substantially altering sensory response dynamics. Choice-related activity in MT can be decomposed in a bottom-up component with the same time-course as the psychophysical kernel, and a top-down component reflecting the decision build-up. These findings suggest that flexible temporal integration can emerge from dynamic interactions between sensory transmission, recurrent decision circuits and temporally structured urgency modulation, providing a mechanistic framework for how neural systems adapt the computation of sensory evidence to changing task demands.

## 1 Introduction

Perceptual decisions rely on the systematic transformation of stochastic sensory signals into action-oriented representations across distributed cortical networks. In motion-direction discrimination paradigms, this computation is mediated by a distinct hierarchical segregation between sensory encoding and decision-related variables. Neurons in the middle-temporal (MT) visual cortex primarily encode sensory motion information, parameterizing physical motion features with directional selectivity [1–3]. Prefrontal and parietal structures such as the lateral intraparietal cortex (LIP) [4–7] and the frontal eye fields (FEF) [8, 9] exhibit sequential ramping activity that scales with the temporal accumulation of evidence toward choice commitment time [10].

Extensive neurophysiological work has characterized the functional dissociation along this sensorimotor axis [11–14], prompting theoretical frameworks to formalize the network mechanics governing information transfer [15]. At the sensory stage, neural firing modulation in area MT is typically modeled as a function of motion direction and coherence through directional tuning curves, where population response amplitudes are scaled linearly or via contrast-gain rectification based on stimulus strength [16, 17]. Downstream decision formation in parietal and prefrontal circuits is formalized either through mechanism relying on sequential sampling of evidence such as drift-diffusion models that implement evidence integration as a continuous stochastic process toward a decision boundary [18–20], or via biophysical network architectures where choice commitment emerges from recurrent, winner-take-all attractor dynamics [21–23]. Rather than executing as fixed feedforward pipelines, these downstream accumulators are highly sensitive to upstream gating. Extensions of such models include dynamic top-down modulations including expected reward evaluations [24], gaze-dependent attentional selection [25], and internal goal-directed drives [26] pre-emptively alter the gain and baseline state of integration circuits prior to evidence onset. This rapid contextual flexibility is mirrored over longer behavioral timescales by structural, experience-dependent plasticity along the sensorimotor axis as agents achieve task mastery [27].

Mechanistically, biophysically realistic recurrent attractor networks propose that slow NMDA-mediated, local recurrent excitation, paired with feedback inhibition within downstream pools, can instantiate the long timescales required for formal temporal integration. This architecture allows downstream decision circuits to smoothly integrate the high-frequency, volatile spike-train inputs transmitted by upstream sensory populations [28]. Crucially, our current modeling framework isolates the fundamental sensorimotor pathway. By explicitly simulating only the sensory encoding layer and the decisional accumulation circuit, we preserve a streamlined, mathematically tractable architecture focused strictly on the canonical principles of evidence-to-intention transformation, including the bidirectional exchange of signals between the two stages.

A central empirical signature of this bidirectional exchange is choice probability (CP), which quantifies the trial-by-trial covariation between a sensory neuron’s response and the animal’s upcoming choice [29]. While CP has traditionally been interpreted as evidence for a feedforward, causal contribution of sensory neurons to behavior [30, 31], accumulating evidence instead points to a substantial top-down component, whereby decision-related signals are fed back to sensory areas after or during evidence accumulation [28, 32]. A recent large-scale meta-analysis across the macaque visual cortex literature reinforces this view, showing that CP magnitude depends systematically on stimulus duration and task structure in ways difficult to reconcile with a purely feedforward account [33]. However, because feedforward and feedback contributions to CP are typically conflated within a single, static task condition, directly dissociating their relative timing and magnitude has remained challenging.

Despite this body of work, how the temporal structure of sensory evidence itself is weighted during accumulation, and how this weighting is instantiated by the interplay between feedforward and feedback signaling along the sensorimotor pathway, remains poorly understood. Here we address this question by training two macaques on a motion-direction discrimination task in which the temporal profile of evidence was systematically manipulated across three conditions, presenting informative motion pulses with either uniform, early-, or late-weighted temporal statistics. Combining psychophysical analyses of choice behavior with population recordings from area MT, we characterize how temporal evidence weighting is expressed jointly at the behavioral and neural levels. We then use a recurrent network model of reciprocally connected sensory and decision populations to test whether condition-specific modulatory signals acting on an otherwise canonical feedforward-feedback architecture are sufficient to reproduce the observed behavioral and neural signatures, and we dissect the distinct contributions of feedforward and feedback coupling to stimulus-driven and choice-related activity. Finally, we use task-trained recurrent neural networks to ask whether temporally structured evidence weighting emerges more generally under variable sensory reliability, independent of the specific biophysical architecture assumed by the reduced circuit model.

## 2 Results

We trained two macaques (subjects N and L) to perform a motion-discrimination task with a highly controlled spatiotemporal stimulus (Fig. 1a). After acquiring fixation, a visual stimulus was presented, consisting of seven consecutive 150 ms motion pulses, without gaps, and subjects reported the net motion direction with a saccade to one of two choice targets. Each pulse consisted of an array of 19 Gabor elements, a subset of which drifted coherently to the left or right while the remaining elements flickered in counterphase, so that the signed motion strength of a pulse, defined as the proportion of coherently drifting elements, was set independently on each pulse. The mean motion strength of each trial was drawn from five levels (strong left, weak left, zero, weak right, strong right). Correct choices were defined by the sign of the cumulative motion strength across the seven pulses, so that every pulse was potentially informative about the reward.

**Fig. 1.**
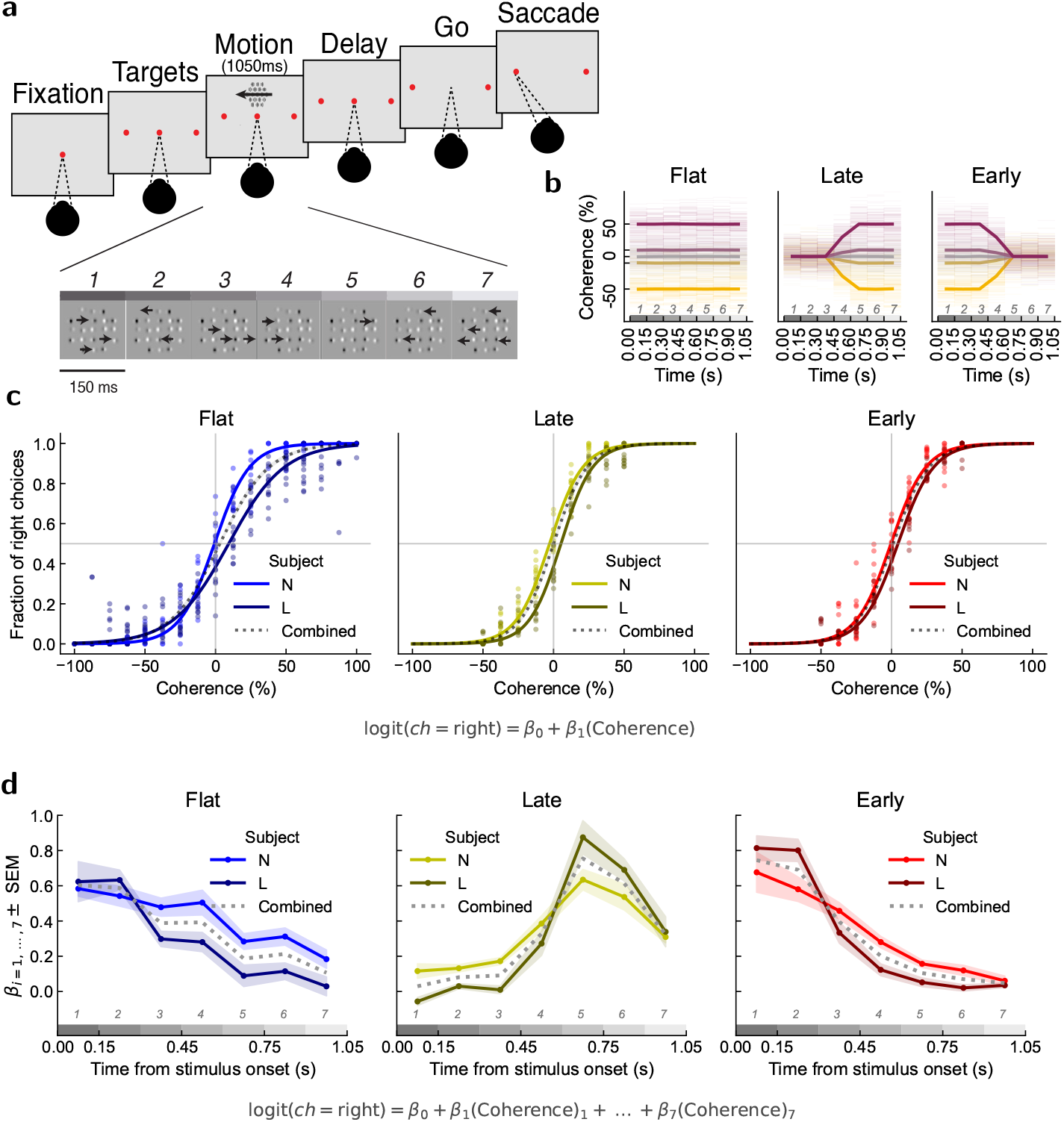
Behavioral paradigm and psychophysical characterization of temporal weighting conditions. **a**. Perceptual motion-discrimination task. Trials start with a fixation point at the center of the screen. Once fixation is acquired, two choice targets appeared on opposite sides. The visual stimuli consist of a sequence of seven consecutive motion pulses, each lasting 150 ms, with independently changing direction and coherence. The signed coherence is given by the proportion of drifting Gabor elements, with positive coherence indicating rightward motion. Following stimulus presentation and a delay period subjects report their choice about the net motion direciton with a saccadic eye movement to a choice target. **b**. Example stimulus time courses showing randomly generated motion strength and coherence across time for the three task conditions (“Flat”, “Late”, “Early”). **c**. Psychometric functions showing the fraction of choices toward right target (positive coherence) versus net motion direction coherence. Dots correspond to average fractions in individual sessions. Solid lines are logistic fits, pooled across sessions for each of the two subjects. **d**. Psychophysical kernels (PK; mean ±SEM across sessions), quantified by the logistic regresion weight [*β*_1_, …, *β*_7_] of each pulse. **c–d**. Gray dotted lines indicate results from fits obtained by pooling trials across sessions and subjects.

We manipulated the temporal statistics of the evidence across three conditions (Fig. 1b). In the flat condition, the expected motion strength was constant across the seven pulses. In the late and early conditions, the expected motion strength was respectively weak at the beginning and strong at the end of the stimulus, or vice versa, so that the informative evidence was concentrated late or early in the trial. Each condition was held fixed within a session and changed across blocks of sessions. This design lets us ask how subjects, and the neural circuits supporting their decisions, weight evidence over time when the statistics of the stimulus change. We first characterize choice behavior, and then examine the neural activity recorded in area MT and its relation to the model.

### 2.1 Effects of temporal weighting of sensory evidence on behavior

Subjects showed consistent performance across conditions (Fig. 1c; Supp. Fig. A1a, Supp. Table A2), although subject N showed more pronounced performance fluctuations in sessions following changes in temporal evidence conditions across blocks (Supp. Fig. A1a). Both subjects exhibited significant sensitivity to the cumulative motion strength across all temporal evidence conditions (Fig. 1c, *β*_1_ in Supp. Table A4). In addition, subject N exhibiting a slight rightward choice bias and subject L a slight leftward choice bias (quantified in terms of *β*_0_ in Supp. Table A4).

Logistic regression of rightward choices as a function of pulse-wise motion strength was used to estimate psychophysical kernels (PKs). Both subjects changed their temporal weighting profiles in the different stimulus conditions, putting generally higher weight to portions of the stimulus with higher expected motion strength (Fig. 1d). Specifically, the flat condition produced sustained PKs throughout stimulus presentation (though earlier pulses are overweighted), whereas the early and late conditions were characterized by larger PK magnitudes during the initial and final pulses, respectively. We further determined the slope of PK profiles by regressing PK = *α*_0_ + *α*_1_**x**, with PK = [*β*_1_, *β*_2_, …, *β*_7_] and **x** = [1, 2, …, 7], which led us to consistently assess negative *α*_1_ across sessions in flat and early conditions, positive *α*_1_ in the late condition (Supp. Fig. A1b).

### 2.2 Temporal weighting and MT activity

The firing rate of MT cells was modulated by the signed motion strength and by the monkey’s choice. In Fig. 2a we pool trials according to average motion strength (strong, weak, zero) and motion direction tuning, and show that the average firing rate follows an approximately linear modulation gradient temporally aligned with motion evidence conditions. The firing rate modulation is sustained across stimulus presentation time for the flat condition, and respectively shows stronger modulation for final or initial pulses in late and early conditions. These firing rate modulation profiles are consistently observed in the two subjects (Supp. Fig. A2).

**Fig. 2.**
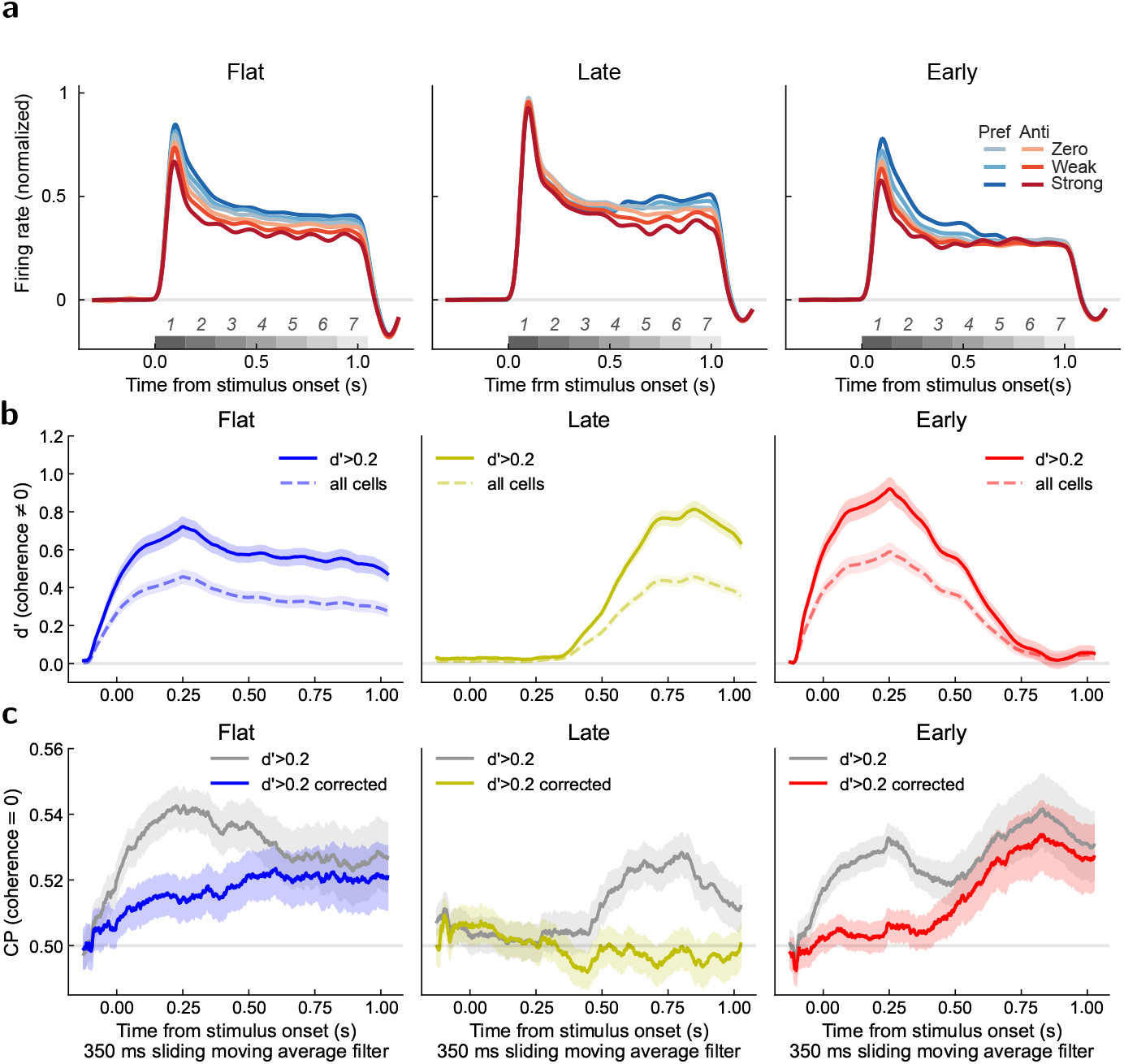
Stimulus and choice selectivity in MT. **a**. Baseline-subtracted, peak-normalized firing rates for the respective cumulative motion strengths (darker for strong, lighter for weak and zero coherence) and choice preference tuning (preferred in blue, or anti-preferred in red) in the three stimulus conditions. Population firing rates were baseline-subtracted by subtracting the mean prestimulus firing rate from each cell’s firing rate and normalized to their peak response across trials. Traces show the mean across cells and trials and were smoothed using a Gaussian kernel (*σ* = 25 ms). Averages are reported across all cells (*n* = 161 Flat, 223 Late, 210 Early), aligning neurons according to their preferred direction (preferred right: *n* = 80 flat, *n* = 108 late, *n* = 82 early; preferred left: *n* = 81 flat, *n* = 115 late, *n* = 128 early). Seperate results for the two subjects are shown in Supp. Fig. A2. **b**. Time-resolved motion discriminability (*d*′ mean± SEM across sessions) for preferred motion direction vs. the opposite direction in the three stimulus conditions. Results are shown for all cells (*n* = 161 Flat, 223 Late, 210 Early) and for cells showing epoch-wise *d*′ *>* 0.2 (*n* = 84 Flat, 87 Late, 116 Early), computed on the time-averaged firing rate during stimulus presentation time (Supp. Table A5). Results for the two subjects are shown separately in Supp. Fig. A3. **c**. Choice Probability (CP mean ±SEM across sessions) for zero pulse-averaged coherence trials across sessions for each of the three temporal weighting conditions. CPs were computed from raw firing rates (gray) and by applying CP correction removing the effect of non-zero motion strength of single pulses on firing rates (colored). Results for all cells and for each subject are shown in Supp. Fig. A4. **b**–**c**. All time-resolved analyes are computed using a moving spike rate window of width *T* = 350 ms sliding with step 1 ms.

We quantified motion discriminability by comparing the firing rate of each cell for opposite motion directions, computing *d*′ between preferred and non-preferred motion directions for trials with mean motion strength between 10% and 50%) across temporal evidence conditions (Fig. 2b). Consistent with the modulation-strength profiles, time-resolved *d*′ remained sustained throughout stimulus presentation in the flat temporal evidence condition, whereas discriminability was preferentially enhanced during the initial or final stimulus pulses in the early and late conditions, respectively. As expected, the magnitude of time-resolved *d*′ was larger when averaging across the subset of highly selective neurons compared to analyses including all recorded units (Fig. 2b), but the temporal profiles remained qualitatively similar. All these time-resolved motion discriminabilty patterns were consistent across subjects (Supp. Fig. A3). Moreover, we computed the pulse-triggered average (PTA) of MT neurons, which revealed stronger responses to initial stimulus pulses with decay over subsequent pulses across all temporal evidence conditions. In addition, the early condition showed a progressive increase in response magnitude toward the final pulses (Supp. Fig. A7).

To examine choice-related activity in MT, We computed choice probability (CP), defined as the probability that a randomly selected zero evidence trial associated with a neuron’s preferred choice elicited a higher firing rate than a randomly selected trial associated with the non-preferred choice [29]. Although the pulse sequences had zero average motion strength across the stimulus period, individual pulses contain coherence values slightly above or below zero. To account for firing-rate fluctuations induced by pulse-specific motion signals, we corrected CP estimates by regressing out motion-induced activity. Figure 2c shows both raw and corrected CPs for neurons with epoch-wise *d*′ *>* 0.2. Note that while the stimuli in these trials are exactly the same across flat, late and early conditions, the CP traces are strikingly different. Raw CP is sustained in the flat and early conditions, with a relatively stable CP profile throughout the trial. In the late condition, raw CP is largely absent in the first half of the trial and emerges only later. Correcting for stimulus flucations selectively removes a large part of CP in early periods in the flat and early condition, and in the late period in the late condition, even though non-zero coherence fluctuations were present throughout the entire stimulus presentation period in all conditions. This decrease in CP is aligned with the periods of high stimulus weight in the evidence weighting profile (psychphysical kernels in Fig. 1d). The corrected, residual CP exhibited a gradual increase over time in the flat and early conditions, whereas CPs in the late condition remained at chance-level, and did not show significant magnitude. We will show how a two-area neural network model can explain this non-trivial patterns of the CP time course through a combination of bottom-up and top-down components.

Notably, we found that the *d*′ of non-zero coherence trials and the CP for zero-coherence trials are significantly correlated in both subjects and across temporal evidence conditions, indicating that motion discriminability and choice-predicting modulation properties overlap to some degree across MT cells (Supp. Fig. A5). In flat and early conditions this correlation was also found for motion-induced corrected CPs, while in late condition as motion-corrected CPs showed chance-level magnitude which did not correlate with the *d*′ selectivity of the recorded cells (Supp. Fig. A6).

### 2.3 The recurrent sensory–decision network model

We modeled perceptual decisions using a recurrent network composed of two pairs of mutually inhibitory neural populations representing sensory and decision-related processing stages (Fig. 3a). The sensory populations capture stimulus encoding mechanisms analogous to those observed in sensory cortical areas such as MT, whereas the decision populations represent downstream circuits involved in evidence accumulation and choice formation, like LIP or FEF. The two stages interact through reciprocal excitatory projections that mediate feedforward (FF) transmission of sensory information and feedback (FB) modulation from decision to sensory populations [28].

**Fig. 3.**
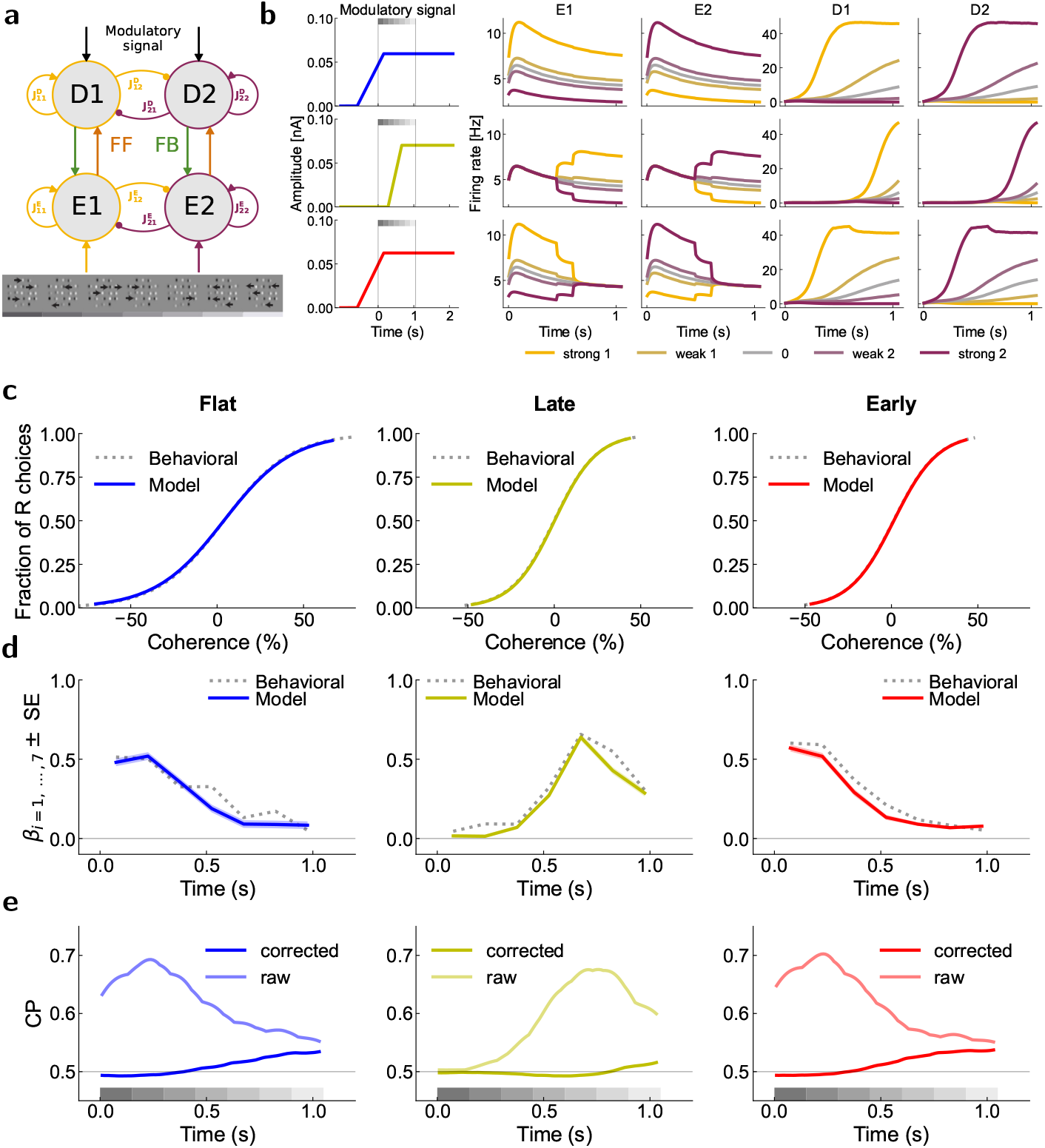
Two-area neural network model and simulation results. **a**. Two-areas model architecture consisting of two pairs of stimulus-selective neural populations, respectively tuned to target 1 or target 2. Populations labelled with E indicate sensory encoding, D indicates decision encoding. Arrows indicate inputs and excitatory connections 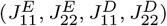, circled edges indicate inhibitory connections 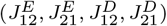. **b**. Modulatory signals and firing rates of the two pairs of populations in the three temporal weighting conditions generated with optimized parameters. From left to right: Modulatory signals, E1, E2 and D1, D2. Top to bottom: flat (blue), late (yellow), early (red). **c**. Logistic fits for choices generated by the two-paired populations model in the three temporal weighting conditions (solid) compared to behavioral performances combined between the two subjects (dotted). **d**. PKs based on simulated choices in the three temporal weighting conditions (solid) compared to behavioral PKs for data combined between the two subjects (dotted). **e**. CP raw (shaded lines) and corrected (solid lines) in sensory population firing rates for the three temporal weighting conditions in zero pulse-average coherence trials. **b–e**. Simulations included 10000 trials for each of the five coherence levels (strong 1, weak 1, 0, weak 2, strong 2).

Sensory populations received stimulus-dependent inputs whose mean amplitude scaled with motion coherence according to the framework of Wong and Wang [22]. In addition, decision populations received modulatory signal inputs implemented as clipped ramp signals characterized by their onset time *t*_0_, duration Δ*t*, and amplitude. Flat and early temporal weighting conditions were captured by sustained modulatory signals, whereas the late condition required delayed modulation (Fig. 3b). These signals were introduced to account for temporal weighting profiles and choice-related activity that could not be reproduced by feedforward sensory integration alone.

Choices were generated by comparing the activity of the decision populations at the end of stimulus presentation. Condition-specific bias terms accounted for the behavioral biases observed experimentally, namely leftward bias in the flat condition, no bias in the late condition, and weak leftward bias in the early condition (Supp. Table A4). A subset of model parameters was fixed (Table 1), whereas the remaining parameters were optimized using a two-stage procedure consisting of an initial grid search followed by Bayesian Adaptive Direct Search (BADS) optimization (Table 2). Parameters shared across the three temporal evidence-weighting conditions were initially set based on values adopted in previous recurrent network models of perceptual decision-making [22], with adjustments to the present model and dataset. In particular, the sensory and recurrent connectivity parameters common to all temporal weighting conditions were manually adjusted, together with initial values of the modulatory input amplitude *I*_*u,e*_, to place the model within a regime that reproduced the characteristic corrected CPs observed in MT neurons. These values served as initial conditions for the subsequent optimization rather than as final parameter estimates. We then performed a grid search to identify suitable parameter ranges and used Bayesian Adaptive Direct Search (BADS) to determine the optimized condition-specific parameters, including the modulatory input amplitude *I*_*u,e*_, onset time *t*_0,*e*_, and duration Δ*t*_*e*_, as well as the subjective choice bias and sensitivity parameters *B*_*e*_ and *σ*_*e*_, for each temporal evidence-weighting condition *e* = flat, late, early. The optimization jointly constrained the model by the observed choices and the PK profiles of temporal evidence weighting. An important consideration is that the sensory population in our model represents an effective node within the broader network contributing to decision formation, rather than a direct reconstruction of the activity measured in MT. The decision-related circuit may receive convergent stimulus information from multiple sensory and extrasensory sources that are not explicitly represented in the present reduced architecture. Consequently, the effective stimulus drive represented by the modeled sensory population may exceed that captured by the activity of the recorded MT population alone. Thus, we did not expect a quantitative correspondence between the absolute magnitude of raw CP in the modeled sensory population and that measured in MT.

**Table 1.**
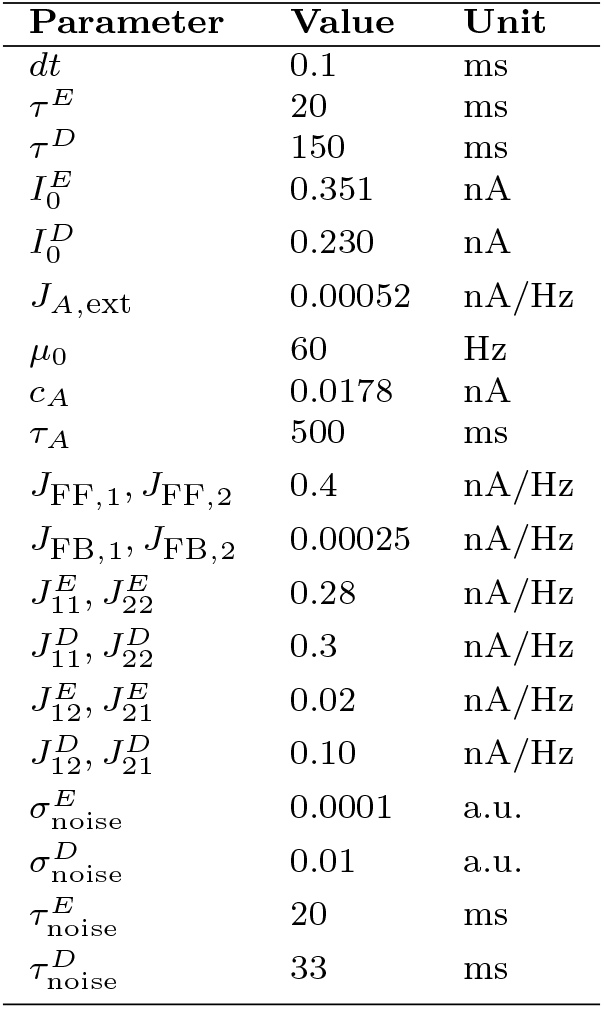
Fixed Parameters manually adjusted to match CP magnitudes in flat condition.

| Parameter | Value | Unit |
| --- | --- | --- |
| $dt$ | 0.1 | ms |
| $\tau^E$ | 20 | ms |
| $\tau^D$ | 150 | ms |
| $I_0^E$ | 0.351 | nA |
| $I_0^D$ | 0.230 | nA |
| $J_{A,\text{ext}}$ | 0.00052 | nA/Hz |
| $\mu_0$ | 60 | Hz |
| $c_A$ | 0.0178 | nA |
| $\tau_A$ | 500 | ms |
| $J_{\text{FF},1}, J_{\text{FF},2}$ | 0.4 | nA/Hz |
| $J_{\text{FB},1}, J_{\text{FB},2}$ | 0.00025 | nA/Hz |
| $J_{11}^E, J_{22}^E$ | 0.28 | nA/Hz |
| $J_{11}^D, J_{22}^D$ | 0.3 | nA/Hz |
| $J_{12}^E, J_{21}^E$ | 0.02 | nA/Hz |
| $J_{12}^D, J_{21}^D$ | 0.10 | nA/Hz |
| $\sigma_{\text{noise}}^E$ | 0.0001 | a.u. |
| $\sigma_{\text{noise}}^D$ | 0.01 | a.u. |
| $\tau_{\text{noise}}^E$ | 20 | ms |
| $\tau_{\text{noise}}^D$ | 33 | ms |

**Table 2.** Simulation parameters specific for temporal weighting conditions optimized via BADS. Tested grid shows parameters used for pre-optimization model evaluation applied coordinate-wise. Underlined values led to identify the best optimization bounds used in random starts of Bayesian adaptive direct search optimization.

| Parameter | Grid-search points | Optimized | Unit |
| --- | --- | --- | --- |
| $I_{u,\text{flat}}$ | 0, 0.02, <u>0.04</u>, <u>0.06</u>, 0.08, 0.1, 0.12 | 0.0596 | nA |
| $I_{u,\text{late}}$ | 0, 0.02, 0.04, <u>0.06</u>, <u>0.08</u>, 0.1, 0.12 | 0.0712 | nA |
| $I_{u,\text{early}}$ | 0, 0.02, 0.04, <u>0.06</u>, <u>0.08</u>, 0.1, 0.12 | 0.0624 | nA |
| $t_{0,\text{flat}}, t_{0,\text{early}}$ | -1.0, -0.8, <u>-0.6</u>, <u>-0.4</u>, -0.2, 0.0 | -588 | ms |
| $t_{0,\text{late}}$ | -0.4, -0.2, 0.0, <u>0.2</u>, <u>0.4</u>, 0.6 | 278 | ms |
| $\Delta t_{\text{flat}}, \Delta t_{\text{early}}$ | 0, 0.2, 0.4, <u>0.6</u>, <u>0.8</u>, 1.0, 1.2 | 742 | ms |
| $\Delta t_{\text{late}}$ | 0, 0.2, <u>0.4</u>, <u>0.6</u>, 0.8, 1.0, 1.2 | 382 | ms |
| $B_{\text{flat}}$ | -0.3, <u>-0.2</u>, <u>-0.1</u>, 0, 0.1, 0.2, 0.3 | -0.120 | a.u. |
| $B_{\text{late}}$ | -0.3, -0.2, -0.1, <u>0</u>, 0.1, 0.2, 0.3 | +0.016 | a.u. |
| $B_{\text{early}}$ | -0.3, -0.2, <u>-0.1</u>, <u>0</u>, 0.1, 0.2, 0.3 | -0.064 | a.u. |
| $\sigma_{\text{flat}}$ | 0, 0.1, 0.2, <u>0.3</u>, <u>0.4</u>, 0.5 | 0.498 | a.u. |
| $\sigma_{\text{late}}$ | 0, 0.1, 0.2, <u>0.3</u>, <u>0.4</u>, 0.5 | 0.531 | a.u. |
| $\sigma_{\text{early}}$ | 0, 0.1, 0.2, 0.3, <u>0.4</u>, <u>0.5</u> | 0.583 | a.u. |

This distinction is particularly relevant when considering the joint constraints imposed by sensory CPs and PKs. Within our network architecture, reducing the stimulus-driven input and recurrent amplification to constrain raw CPs to the magnitude observed in MT substantially decreases the stimulus-dependent signal propagated to the decision populations, resulting in poorer choice estimation likelihood and on PKs that are markedly smaller than those observed empirically in MT (Supp. Fig. A9). Conversely, the effective stimulus drive required to reproduce the experimentally observed magnitude and temporal structure of the PKs necessarily produces larger raw CPs in the modeled sensory population. We therefore constrained the baseline neural dynamics based on the magnitude of corrected CPs, while optimizing the condition-specific modulatory parameters to jointly reproduce psychometric performance, PK amplitude and temporal structure, and condition-dependent choice-related activity.

Once the baseline neural dynamics parameters were fixed, we used a two-stage optimization procedure to determine the condition-specific parameters in the three temporal evidence weigthing configurations. To identify suitable parameter ranges, we first run a coarse grid-search across condition-specific parameters, then we proceeded to a finer optimization with a simulation-based inference approach using Bayesian Adaptive Direct Search (BADS). Specifically, for each condition *e* = flat, late, early, we optimized the modulatory signal amplitude *I*_*u,e*_, onset time *t*_0,*e*_, and duration Δ*t*_*e*_, together with the subjective choice bias and sensitivity parameters *B*_*e*_ and *σ*_*e*_. The optimization jointly maximized the likelihood of observed choices and minimized discrepancies between empirical and simulated PKs (Methods). The fixed parameters are reported in Table 1, while the BADS-optimized parameters are reported in Table 2.

The optimized model reproduced behavioral signatures across temporal weighting conditions. Simulated psychometric functions closely matched behavioral performance across stimulus coherences (Fig. 3c), whereas simulated PKs captured the experimentally observed temporal weighting profiles (Fig. 3d). In addition, we tracked choice-selective activity profiles defined as the difference in firing rates between chosen and unchosen trials in sensory units for zero pulse-average coherence trials across temporal evidence conditions in (Fig. 3e). Simulated sensory activity also reproduced the temporal structure of the PTA computed on MT neurons activity (Supp. Fig. A8).

### 2.4 The role of the respective parameters

To examine the role of feedforward signaling, we varied the FF coupling strength (*J*_FF_). Reducing *J*_FF_ progressively impaired behavioral performance, reaching chance-level choice accuracy as it approached zero (Supp. Fig. A10). Perturbing FF strength strongly affected temporal evidence-weighting profiles, as shown by PKs. Weak FF coupling produced near-zero PKs, intermediate values generated larger PK amplitudes, whereas strong FF coupling led to increasingly skewed PKs toward the beginning of the stimulus presentation (Supp. Fig. A10). Increasing FF coupling also progressively accelerated and amplified choice-selective differences between neural populations, as captured by the CPs. Weak FF coupling produced little choice-selective separation, whereas stronger coupling generated earlier and larger differences in decision-circuit activity, resulting in increased raw CPs (Supp. Fig. A10). Notably, stronger FF coupling also increased corrected CPs. This effect arises from the stronger influence of stimulus-driven activity on the decision circuit: although CP correction removes the component of neural activity that can be linearly explained by stimulus motion coherence, stimulus information is transformed and integrated nonlinearly before the decision is formed. Consequently, increasing *J*_FF_ could also indirectly increase corrected CPs, although its largest effect was on raw CPs. Despite these pronounced effects on behavior and choice-related activity, FF perturbations had no substantial influence on sensory PTA profiles (Supp. Fig. A10).

In contrast, varying FB coupling (*J*_FB_) produced relatively small changes in psychometric performance and PK structure. Instead, increasing FB strength selectively enhanced choice-related activity, selectively increasing the neural response to preferred targets, thus mainly increasing the latest part of both raw and corrected CPs, with major influence on corrected CPs (Supp. Fig. A11). Stronger feedback also directly led to increase the magnitude of late sensory PTA (Supp. Fig. A11). Both effects are consistent with a choice-selective top-down modulatory effect.

Together, these results indicate that FF interactions primarily shape stimulus-driven sensory response modulation as shown by raw CPs, whereas FB interactions mainly contribute to late choice-related signalling captured by corrected CPs, consistent with results observed in MT (Fig. 2c) and with previous modeling work dissociating early FF and late FB contributions to CP in neural dynamics [28].

We next dissected the role of the modulatory signals *I*_mod_. Varying the modulation onset time (*t*_0,*e*_) across conditions (*e* = {flat, late, early}) revealed strong condition-specific effects. Delaying modulation signal onset substantially impaired behavioral performance in the flat and early conditions, whereas the late condition exhibited a narrow optimum near the fitted value, with both earlier and later onsets drastically reducing choice performance (Supp. Fig. A12). Manipulating modulation onset time systematically shifted PK profiles, with early onsets emphasizing initial stimulus information and delayed onsets increasing the influence of late evidence, primarily as a consequence of shifts in the dynamics of decision units activity (Supp. Fig. A12). The PTA profiles remained largely unaffected by these manipulations (Supp. Fig. A12).

We then manipulated the slope of the clipped ramp signals by scaling their duration parameters (Δ*t*_*e*_) by a factor *k* ∈ [0, 2]. Altering ramp slopes had little effect on psychometric performance (Supp. Fig. A13) but systematically modulated the temporal structure of PKs and choice-selective activity. Increasing *k* enhanced late weighting and delayed choice-specific modulation, whereas smaller values shifted both effects toward earlier stimulus periods (Supp. Fig. A13), deriving from changes in the associated modulation signals and decision circuit responses (Supp. Fig. A13). Changing the slope did not affect the PTA profiles (Supp. Fig. A13).

Lastly, we examined the influence of the condition-specific subjective choice bias term. Varying the subjective bias *B*_*e*_ across conditions *e* ∈ flat, late, early shifted the psychometric functions and show that the optimizing bias values leads us to finely reproduce the subjective asymmetries observed experimentally (Supp. Fig. A14). We also tested the effects of changing self-excitatory strengths in sensory 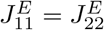 (Supp. Fig. A15) or decision circuits 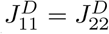 (Supp. Fig. A17), the cross-inhibitory strengths in sensory sensory 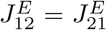 (Supp. Fig. A16)) or decision circuits 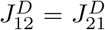 (Supp. Fig. A18), and the effects of noise in the sensory 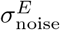 (Supp. Fig. A19) or in the decision units 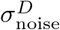 (Supp. Fig. A20).

### 2.5 Optimality of non-uniform temporal weighting in task-trained RNNs

Because reward in our task depended on the sign of the sum of the seven motion pulses, a noise-free, optimal, observer would weight all pulses equally, so the primacy and recency kernels of the animals (Fig. 1d) appear suboptimal. We asked whether non-uniform weighting can instead be the optimal solution when evidence integration is corrupted by noise (either sensory or internal noise). To test this without assuming any particular circuit mechanism, we trained recurrent neural networks (RNNs) by supervised learning to maximize accuracy, which is the ideal-observer objective for this task, and examined the temporal weighting they converged to. Any non-uniform kernel in these networks can therefore only arise from the interaction between the stimulus statistics and the noise, not from a built-in bias, a modulatory signal or a cost on integration.

Networks (256 ReLU units, *τ* = 100 ms) received a fixation input and a stream of seven noisy motion pulses (150 ms bins) generated with the flat, late or early statistics of the task, and reported the sign of the net motion at the end of the trial. We trained separate networks on each condition at each of six noise levels and estimated psychophysical kernels (PKs) with the same logistic regression used for analyzing the behavioral data. Note that these independetly trained RNNs are not suitable to identify the circuit mechanism underlying flexible temporal integration, which was the subject of the circuit model above.

We found that networks learn all three conditions and their kernels follow the stimulus statistics (Fig. 4). At high noise (*σ*_*S*_ = 3), networks trained on the flat condition weighted all pulses equally, networks trained on the late condition showed a strong recency kernel, and networks trained on the early condition showed a strong primacy kernel (Fig. 4). These kernels resemble the patterns of late and early PKs of the animals (Fig. 1d). At low noise, kernels were nearly uniform in all conditions and psychometric curves were steep (Fig. 4a,b). Increasing noise reduced overall kernel magnitude, giving rise to a flattened psychometric function, and progressively shifted the kernels in the late and early conditions, while the flat-condition slope stayed near zero (Fig. 4c).

**Fig. 4.**
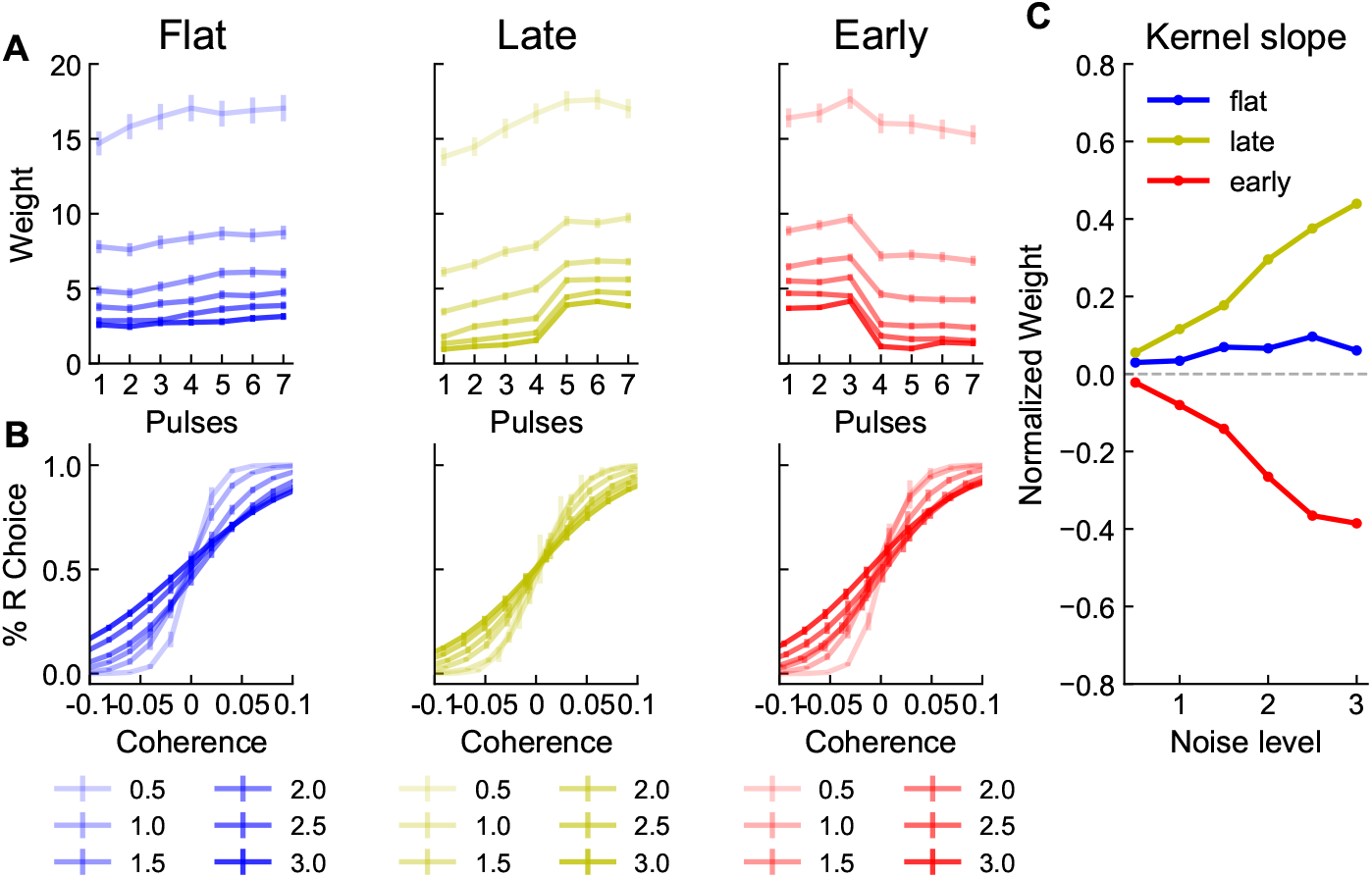
Non-uniform temporal weighting emerges as an optimal strategy in task-trained RNNs. RNNs were trained independently on the flat, late and early conditions at noise levels *σ*_*S*_ = [0.5, 3] (light to dark colors). Curves show the mean across 10 networks per condition and noise level. **a**. Psychophysical kernels (logistic regression weights of the seven pulses). **b**. Psychometric functions (fraction of right choices vs. motion strength). **c**. PK slope as a function of noise level (positive values indicate recency, negative values primacy and zero uniform weighting).

Taken together, this shows that non-uniform integration is not necessarily evidence of suboptimal processing. When evidence is unreliable or corrupted by internal noise, the best strategy is to emphasize the periods in which the expected signal is strongest, and uniform weighting becomes optimal only when noise is negligible. The RNNs thus indicate that the primacy and recency kernels of the animals are compatible with an accuracy-maximizing strategy under noise in the early and late conditions. In the flat condition, the monkeys primacy kernels are actually suboptimal.

## 3 Discussion

We combined behavioral, neurophysiological, and computational approaches to characterize how the temporal structure of sensory evidence shapes perceptual decision making. Manipulating whether informative motion pulses were presented uniformly, or concentrated early or late within a trial, produced systematic shifts in psychophysical kernels that closely tracked the temporal evidence-weighting condition, indicating that subjects flexibly adjusted how much weight each moment of evidence received rather than integrating all pulses uniformly [13, 14]. Extending previous work [13, 14], we developed a circuit-level model that explains behavioral and neural signatures of flexible decision making, and a normative account of why the psychophysical kernels take the shapes they do. Consistent with representing instantaneous motion information, we found that motion discriminability (*d*′) in MT tracked the temporal profile of task-relevant evidence. Choice-predictive modulation (choice probability, CP) could be decomposed in a stimulus-driven component, and a later, gradually rising component. The correlation we observed between *d*′ and CP across neurons and conditions is consistent with the view that sensory and choice-related signals are not fully segregated processing stages but have at least partially overlapping representations [29, 32].

Our two-population recurrent network model, built on a reciprocally connected sensory-decision architecture [28], reproduced both the behavioral psychometric functions and psychophysical kernels across all three temporal weighting conditions using only condition-specific modulatory ramp signals. Critically, dissecting the roles of feedforward and feedback coupling recovered a dissociation consistent with prior modeling and empirical work [28]: feedforward strength primarily governed the magnitude and timing of raw choice probability and the overall reliability of the psychophysical kernel, whereas feedback strength selectively sculpted late choice-related activity and corrected CP, without substantially altering behavioral sensitivity. This double dissociation supports the interpretation that top-down, choice-related signals are conveyed to sensory areas largely independently of the feedforward pathway that carries stimulus information upstream, in line with broader proposals that decision formation reflects a continuous, bidirectional exchange between sensory and decision-related circuits rather than a strictly feedforward accumulation process [11, 12, 15].

RNNs trained on the same task show that non-uniform temporal weighting of evidence does not always indicate a departure from optimal integration. If evidence were observed without noise, the reward-maximizing strategy in this task would be to weight all pulses equally. Networks trained only to maximize accuracy did so at low noise, but at higher noise they converged on kernels that follow the temporal profile of the signal, with recency in the late condition and primacy in the early condition. The animals’ kernels in these two conditions are therefore compatible with an accuracy-maximizing strategy, rather than evidence of suboptimal integration. It should be noted that the networks were trained separately for each condition and therefore say nothing about how the animals acquired their strategies. The slow drift of kernel slopes over sessions after each change of condition (Supplementary Fig. A1b) suggests gradual learning rather than trial-by-trial switching, so “flexible” should be understood on the timescale of days. The flat condition is a clear exception to optimality: both animals weighted early pulses more than late ones, which is not accuracy-maximizing. Plausible explanations include bounded integration or early commitment [34], or a default strategy learned under natural statistics.

Together, these results argue that flexible temporal weighting of evidence is implemented through condition-specific modulatory signals acting on an otherwise canonical sensory-to-decision pathway [25, 26], and that feedback projections from decision to sensory areas play a specific role in shaping late choice-related activity independently of overall task performance. Several open questions remain. First, our model treats the sensory population as an effective node summarizing input to the decision circuit rather than a literal reconstruction of MT, and future work relating model and neural CP magnitudes more quantitatively will require explicitly modeling the convergence of multiple upstream sources [35–37]. Second, the origin of the modulatory ramp signals themselves remains unspecified in the current architecture; candidate sources include value-based, attentional, or goal-directed modulation shown to shape decision circuits in other contexts [38–41], potentially routed through the laminar- and frequency-specific circuits implicated in top-down control [42, 43]. Addressing these questions will help clarify how the brain flexibly reallocates decision weight across time in response to task demands.

## 4 Methods

### 4.1 Experimental settings and reproducibility

Stimuli were generated in MATLAB (MathWorks) using the Psychophysics Toolbox for visual presentation control[44], with precise event timing and registration handled via a Datapixx I/O system (Vpixx)[45]. Eye position was tracked using an Eyelink eye tracker (SR Research), sampled at 1 kHz. Subject L was seated 57 cm away from a 150× 86 cm rearprojection screen (IRUS; Draper) covering the central 106 ×73 degrees of visual angle. Images were projected onto the screen by a PROPixx Projector (VPixx Technologies) driven at a resolution of 1920 ×1080 pixels at 120 Hz. Subject N viewed stimuli on a 121.8× 68.5 cm LCD (LG) display (resolution 1920× 1080 pixels; refresh rate 60 Hz; background luminance = 26.49 cd/m^2^, corrected to have a linear g function) from a distance of 118 cm (such that the screen width subtended 54 degrees of visual angle). Auditory feedback was played at the end of every trial, and fluid reward was delivered through a computer-controlled solenoid valve.

### 4.2 Experimental procedures and neural recordings

Two adult rhesus macaques served as subjects (referred to as N and L, respectively, hereafter), aged 10 and 6 years, female and male, respectively. All procedures were performed in accordance with US National Institutes of Health guidelines, were approved by The University of Texas at Austin Institutional Animal Care and Use Committee, detailed in previous work[46]. Extracellular recordings were accomplished using single electrodes (glass-coated tungsten; Alpha Omega) and multisite linear-electrode arrays (U-probe or V-probe, Plexon; 50–150 *µ*m spacing). Anatomically, MT area was targeted using cranial landmarks or structural MRI, electrode depths, sulcal anatomy (gray/white boundaries) and functional mapping. Functionally, MT was identified based on receptive field size and a preponderance of directionally selective cells[1].

Recording from microelectrode arrays yielded a wide sample of units (*n* = 119 in subject N, 475 in subject L, 594 in total). For analyses of stimulus selectivity, we included all units and separated the ones that were sensitive to the net direction of motion during stimulus presentation, identifying units with *d*′ *>* 0.2 (Supp. Table A5).

### 4.3 Behavioral paradigm

Subjects were trained to perform a motion discrimination task via saccadic report to one of two targets located on opposite sides of the screen. Each trial began with fixation of a central point for 750–1300 ms, followed by the appearance of two choice targets, which remained visible throughout the trial. After a 500–1000 ms delay, a motion stimulus consisting of seven consecutive 150-ms pulses was presented without inter-pulse gaps (9 frames at 60 Hz; 18 frames at 120 Hz). Following stimulus offset, an additional 500–1000 ms delay was imposed, after which the fixation point disappeared, serving as a “choice-go” cue instructing subjects to initiate a saccadic report within 600 ms. Trials were aborted if fixation was broken before the go cue (exceeding 3° around fixation), and responses were considered valid if saccade endpoints landed within 3° of either target. Subjects were rewarded for selecting the target consistent with the net average motion across the seven pulses. The task timeline is illustrated in Fig. 1a.

The stimulus consisted of a hexagonal grid (5–7° across, scaled by eccentricity) of 19 Gabor patches (0.9 cyc/deg). Spatial frequency was chosen to match the peak selectivity of MT neurons[47]. Inter-element spacing and Gabor size were scaled to 10% of eccentricity, approximating average V1 receptive-field size and preventing overlap[48]. Each motion pulse contained a subset of Gabor elements assigned coherent motion, while the remaining elements underwent counterphase flicker noise. The motion signal and noise elements were randomly assigned within the spatial array on each pulse. The number of signal elements was determined as a rounded integer drawn from a Gaussian distribution, *X*_*i*_ = round(*Z*_*i*_), with *Z*_*i*_ ~ *N* (*µ*_*k*_, *ν*), for *i* = 1, …, 7. Thus, *µ*_*k*_ denotes the mean across pulses corresponding to *k* = 1, …, 5 trial types, designed to yield mean coherence levels of {-50% “strong left”, −10% “weak left”, 0% “no coherence”, 10% “weak right”, 50% “strong right”}, and *ν* corresponded to a standard deviation across pulses of 5% for all five levels. Each pulse therefore contained a stochastic number of coherent Gabor elements, while the expected value across pulses was fixed to the predetermined mean *µ*_*k*_. Subjects were rewarded for selecting the target consistent with the sign of the motion pulse sequence sum (i.e., left if the net motion direction was negative, right if positive), independent of the mean *µ*_*k*_ of the generating distribution. In zero-coherence trials, reward was assigned randomly with probability 0.5.

Both subjects performed the task under three configurations of temporal stimulus statistics [13], illustrated in Fig. 1b. In the flat condition, expected motion strength was constant across pulses, i.e. *µ*_*k*_ was fixed for all *i* ∈ [1, 7]. In the late condition, motion strength was reduced in early pulses and increased in later pulses, such that *µ*_*k*_ was set to 0 at *i* = 1 and reached its full value by *i* = 7 (Fig. 1b). In the early condition, the reverse temporal profile was used, with full motion strength at early pulses decaying toward 0 at later pulses (Fig. 1b). In the late and early conditions, this transition was implemented via sigmoidal modulation with midpoint *i* = 4 and slope 0.3, yielding average pulse-wise motion strengths {-25%, −5%, 0%, 5%, 25%}. Temporal configuration was fixed within sessions and changed across blocks of consecutive sessions with same condition (Supp. Fig. A1, Supp. Table A1). The number of trials available per condition and the fractions of correct trials are reported in Supp. Table A2, whereas trials per coherence level are reported in Supp. Table A3.

### 4.4 Behavioral data analyses

We track the fraction of choices for right target (positive net motion direction) in pulse-averaged coherence bins, and fitted the choice probability to the logistic model:

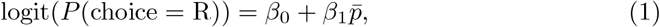

where 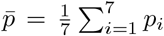 is the average pulse motion coherence within each experimental trial. Statistical significance of the logistic regression coefficients was assessed using Wald tests, and two-sided *p*-values for the estimated coefficients were obtained from the fitted model (Supp. Table A4). We derived Psychophysical Kernels as PK = [*β*_1_, …, *β*_7_] quantifying the logistic weight of the *i*th pulse on rightward choice probability via the regression:

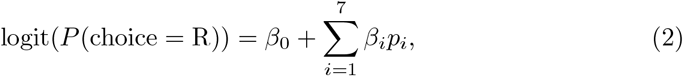

with *p*_*i*_ indicating the motion coherence level of the *i*th pulse. Statistical significance of model weights was assessed using Wald tests based on the asymptotic normality of model estimates. Two-sided *p*-values were obtained under a standard normal distribution approximation.

### 4.5 Neural data analyses

#### 4.5.1 Motion selectivity (*d*′)

We quantified motion direction selectivity using the discriminability index

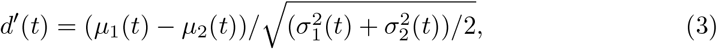

where *µ*_1_(*t*) and *µ*_2_(*t*) are the mean firing rates at time *t* for trials corresponding to the two opposite motion directions in non-zero coherence (strong or weak) trials, 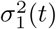 and 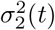 are the respective variances across trials. Time-resolved *d*′ traces were computed using 350 ms sliding windows, starting at *t* = −300 ms, with 1 ms step resolution. For neurons exhibiting negative choice tuning (i.e., higher firing rates for the non-preferred choice direction), *d*′ values were set to −*d*′ suc that the overall *d*′ resulting from cell-averages consistently reflected sensitivity aligned to preferred choice tuning.

#### 4.5.2 Choice Probability

Choice probability (CP) quantified the area under the receiver operating characteristic (ROC) curve (AUC), providing a measure of how well trial-by-trial firing-rate fluctuations predicted the behavioral choices[29]. For each recorded unit, trials were separated according to behavioral choice (target left vs. target right). Considering a neuron exhibiting higher firing *x*(*t*) at time *t* for rightward choices, a decision threshold *θ* was applied to the firing rate such that responses exceeding threshold predicted a rightward choice, whereas responses below threshold predicted a leftward choice. For each threshold value, true-positive and false-positive rates were computed as TPR = TP*/*(TP + FN) and FPR = FP*/*(FP + TN), respectively, where TP, FP, TN, and FN denote true positives, false positives, true negatives, and false negatives. Sweeping *θ* across the full range of firing rates values generated the ROC curve by tracing TPR against FPR. CP was then defined as the area under ROC curve, corresponding to the probability that a randomly selected trial associated with the neuron’s preferred choice exhibited a higher firing rate than a randomly selected trial associated with the non-preferred choice. Time-resolved AUC traces were computed using the same 350 ms sliding-window on the firing rates, as in the *d*′ analyses. For neurons exhibiting negative choice tuning (i.e., higher firing rates for the leftward choice direction), AUC values were transformed as 1− AUC such that population-averaged CP values consistently reflected predictive strength for the rightward choice direction.

To correct CP estimates for non-zero fluctuations under zero-average coherence conditions, we removed the component of neural activity attributable to motion-induced changes in firing rate, following [32]. Let *x*(*t*) denote the raw firing rate in time bin *t*. We estimated the firing rate expected from the motion stimulus alone by fitting a linear regression model, 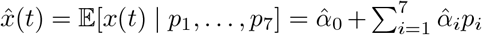, where *p*_*i*_ denotes the motion strength at the *i*th pulse. The residual firing rate, 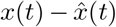, was used to compute corrected CP via AUC traces following TPR and FPR definitions. In both raw and corrected versions, CP is referenced to estimate rigthward target selection by combining the two simulated firing rates as CP = AUC(*x*^*E*^)*/*2 + (1 − AUC(*x*^*E*^))*/*2.

#### 4.5.3 Pulse-Triggered Average (PTA)

The pulse-triggered average (PTA) quantifies the change in spike rate evoked by a stimulus pulse of nominally unitary strength [12]. Spike counts and pulse amplitudes were binned at 10 ms resolution. We denote with *p*(*t*) the motion coherence level of the pulse stimulus at time *t*, and *x*(*t*) the respective spike rate. Trials were concatenated so that the stimulus vector **p** has length *T*, the total number of time bins, and is zero everywhere except at pulse onset times. For a pulse at time *t*, we reference *p*(*t*) *>* 0 for pulses in the preferred direction and *p*(*t*) *<* 0 for pulses in the anti-preferred direction. We compute the temporal response of each neuron by constructing lagged stimulus design matrices. For pulse *k*, the matrix **D**_*k*_ ∈ ℝ^*T* ×*L*^ contains *L* time-lagged copies of the stimulus vector **p** aligned to that pulse, with zeros elsewhere. The full design matrix for all *K* = 7 pulses is **D** = [**D**_1_, **D**_2_, …, **D**_*K*_] ∈ ℝ^*T* ×(*KL*)^. At this point, the mean-subtracted spike rate matrix 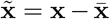 is projected onto **D** using ordinary least squares: 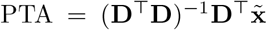. This formulation accounts for correlations across pulses and lags and allows inclusion of all trials, increasing statistical power. The resulting matrix is PTA ∈ ℝ^*L*×*K*^, with each column representing the temporal kernel of a single pulse. In our settings, we use *L* = 60, covering 600 ms after each pulse.

### 4.6 Neural network models

#### 4.6.1 Sensory Circuit

The synaptic gating variable 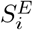 represents the fraction of open excitatory synapses for neuron *i* in the sensory area. Its dynamics are given by

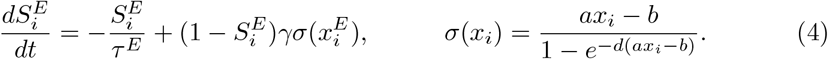

Here, *τ* ^*E*^ is the excitatory synaptic time constant, and *γ* = 0.641 scales the contribution of the neuron’s firing-rate transfer function *σ*(*x*_*i*_). The total synaptic input *x*_*i*_ to unit *i* = {1, 2} is multiplied by the synaptic gain *a* = 270 Hz/nA, shifted by the firing threshold *b* = 108 Hz, and shaped by the temporal scaling factor *d* = 154 ms.

The total input to each sensory neuron combines recurrent excitation, cross-coupling inhibition, feedback, baseline input, stimulus, and noise:

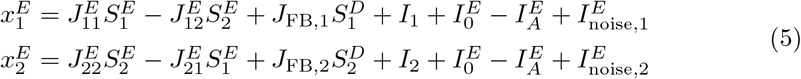

*J*_*ii*_, *J*_*ij*_, and *J*_FB,*i*_ are the recurrent, cross-coupling, and feedback weights.

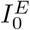 represents background input incorporating adaptation from MT neurons, and

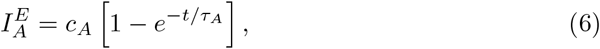

is the adaptation term, with amplitude *c*_*A*_ and effective temporal decay factor *τ*_*A*_. The noisy stochastic fluctuations are modeled as an Ornstein-Uhlenbeck processes:

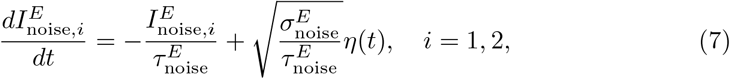

where *η*(*t*) is a white Gaussian process with zero mean and unit variance, 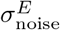 is the 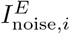 noise variance and 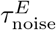 is the synaptic decay time.

The external input current *I*_*i*_ is modulated over time:

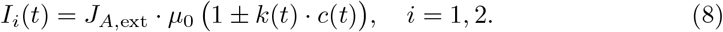

In the early condition, we set *k*(*t*) = 0.4*t*^2^ 0.12*t* + 1 to recover the u-shaped time-dependent scaling of the sensory responses observed in MT data (Supp. Fig. A8). For flat and late conditions, we set *k*(*t*) = 1 without additional shaping.

#### 4.6.2 Decision Circuit

Decision neurons follow the same synaptic gating dynamics as sensory neurons:

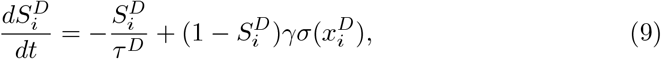

and we adopt the same *γ* = 0.641 and the same transfer function *σ*(*x*_*i*_), *i* = {1, 2} as in the sensory units. The total input combines recurrent excitation, cross-coupling inhibition, feed-forward input, baseline input, modulation signal, and noise:

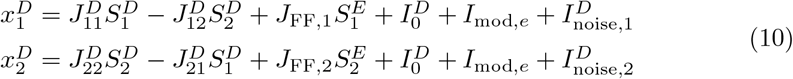

*J*_*ii*_, *J*_*ij*_, and *J*_FF,*i*_: recurrent, cross-coupling, and feed-forward weights, *i, j* = [1, 2].

Stochastic fluctuations in the decision neurons are Ornstein-Uhlenbeck processes:

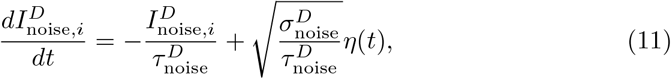

with *η*(*t*) a zero-mean and unitary variance white Gaussian noise process, 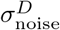 is the 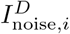 noise variance and 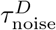 is the synaptic decay time.

The task-dependent modulatory signal *I*_mod,*e*_ is defined for *e* = {flat, late, early}:

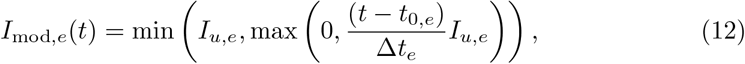

as a clipped ramp starting at *t*_0,*e*_ and with slope Δ*t*_*e*_, and maximal amplitude *I*_*u,e*_.

#### 4.6.3 Choice Estimation

Choice probabilities were modeled using a logistic function of the decision variable derived from normalized decision-population activity, i.e. by assigning the choice to the firing response of the dominant decision unit.

We define a decision variable on firing rates of both decision units at *T* = 1.05 s:

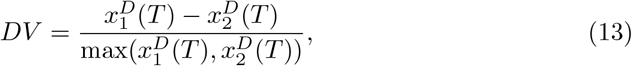

ranging [− 1, +1] in all temporal evidence conditions and coherence levels (strong, weak, 0). For each trial, the *DV* was transformed into a choice probability via:

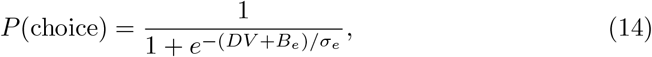

including a bias term *B*_*e*_, and a sensitivity factor *σ*_*e*_, *e* = {flat, late, early} controlling the shape of the psychometric curve, accounting for subjective biases, most pronounced and significant in flat and early conditions (Fig. 1c, Supp. Table A4).

Binary choices were generated by comparing *P* (choice) to a uniformly distributed *u* ~ *U*(0, 1), setting *ch* = 0 (left) when *u < P* (choice), *ch* = 1 (right) otherwise.

#### 4.6.4 Parameters Optimization

We first adapted the baseline network in flat temporal evidence weighting condition, starting from parameters in the original Wong–Wang framework and subsequent extensions [22, 28], adjusted to obtain a regime in which the model reproduced physiologically realistic CPs computed on MT activity. This initial stage determined the parameters governing the main intrinsic dynamics of the network, including effective decay variables *τ*_*E*_ and *τ*_*D*_, background currents 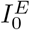 and 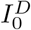, stimulus coherence strengths *J*_*A*,ext_, *µ*_0_, adaptation parameters *c*_*A*_ and *τ*_*A*_, feedforward and feedback coupling strengths *J*_FF_ and *J*_FB_, recurrent connectivity weights 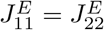 and 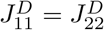, mutual inhibitory weights 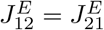 and 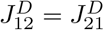, and noise parameters 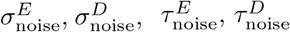,. All these were subsequently held fixed at the values reported in Table 1.

The remainder of the model parameters were then optimized to reproduce the observed behavioral effects of temporal evidence weighting, across flat, early, and late conditions. We used simulation-based inference with a likelihood-based objective function [49, 50]. The optimized parameters comprised condition-specific modulation amplitudes (*I*_*u,e*_), modulation onset times (*t*_0,*e*_), modulation durations (Δ*t*_*e*_), and subjective choice parameters (*B*_*e*_ and *σ*_*e*_), for the respective *e* ∈ {flat, late, early}.

We first performed a coarse, coordinate-wise, grid search over predefined parameter ranges to identify regions of the parameter space providing a good correspondence with the behavioral data (Table 2). These candidate regions were subsequently refined using Bayesian Adaptive Direct Search (BADS), implemented in PyBADS [51, 52]. This approach helped us identify parameter boundaries for the BADS optimization, thus to identify optimized values based that best reproduced behavioral readout (Table 2). For each optimization run, parameter values were initialized by uniform sampling within the corresponding candidate-region bounds and iteratively updated by the BADS algorithm. For every candidate parameter set, the network simulated 1500 trials at each of the five coherence levels, yielding a total of 7500 simulated trials. We initialized the optimization across 5 independently sampled random starting points. Model-generated choices were used to estimate psychometric functions and psychophysical kernels, computed using the same logistic-regression procedure applied to the behavioral data, for each temporal-weighting condition. Model fit quality was evaluated using a composite loss function. First, we computed the negative log-likelihood (NLL) of the simulated data quantifying the accuracy of model-generated choices estimations. Second, we quantified the standard error (SE) between simulated and empirical psychophysical kernels across pulse-specific regression weights.

The optimization of parameters minimized the composite loss function combining NLL and SE: ℒ = Σ_*e* ∈ {flat,late,early}_ (NLL_*e*_ + *λ* SE_*e*_), with *λ* = 10^3^ chosen to balance the numerical scales of the two terms across conditions (NLL ~ 10^4^ and SE ~ 10^−1^).

### 4.7 Recurrent Neural Network Simulations

To investigate how the temporal structure of sensory evidence influences decision-making strategies, we trained recurrent neural networks on a pulse-based perceptual decision-making task implemented in the NeuroGym framework [53, 54]. Networks received sequential streams of noisy sensory evidence and were trained to classify the sign of the accumulated evidence over each trial.

The model consisted of a continuous-time recurrent neural network implemented in PyTorch [55]. Hidden-state dynamics were governed by:

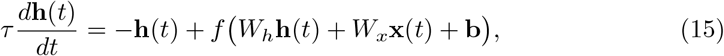

where **x**(*t*) denotes the sensory input, **h**(*t*) the hidden state, *W*_*x*_ and *W*_*h*_ are input and recurrent weight matrices, **b** is a bias term, and *f* (·) is a rectified linear unit (ReLU) nonlinearity. Simulations were performed with a membrane time constant of *τ* = 100 ms and a discrete update interval of Δ*t* = 150 ms. The recurrent layer contained 256 hidden units, and a linear readout layer projected hidden activity onto two decision outputs corresponding to leftward and rightward choices.

Stimuli consisted of seven sequential evidence pulses presented during a trial. For each trial, a coherence value was sampled from the set {−50%, −20%, 0%, 20%, 50%}. Evidence was distributed across time according to different temporal profiles, ranging from uniform average evidence throughout the stimulus period, to profiles in which informative evidence was concentrated at the beginning or at the end of the trial, respectively following flat, early and late temporal weighting conditions. Importantly, coherence magnitudes were identical across conditions, allowing the temporal distribution of evidence to be manipulated independently of overall stimulus strength. To examine how sensory reliability influences evidence integration, we added Gaussian noise to the motion stimuli. Input noise levels ranged from *σ*_*S*_ = 0.5 to 3.0 in increments of 0.5. Increasing noise reduced the reliability of individual evidence pulses while leaving the underlying stimulus statistics unchanged, enabling us to assess how temporal weighting strategies and network dynamics depend on sensory uncertainty. The correct response on each trial was determined by the sign of the mean evidence across all pulses. Trials with exactly zero mean evidence were assigned a random target label. Networks were trained using supervised learning with online trial generation from the NeuroGym environment. At each optimization step, minibatches of 64 trials were sampled, and network parameters were optimized using the Adam optimizer with a learning rate of 10^−3^. Training lasted for 6000 optimization steps. We assessed each combination of temporal evidence profile and input-noise level over 10 trained networks starting from independent random initializations.

Following training, each network was evaluated on 15000 previously unseen trials generated from the same task distribution. Choices were determined by the output unit with maximal activity at the final decision time step. As in behavioral analyses, psychometric functions were obtained by computing the proportion of correct motion detection choices as a function of stimulus coherence and fitting sigmoidal functions to the resulting choice probabilities. Temporal weighting strategies were quantified using psychophysical kernels estimated through logistic regression of network choices onto pulse position. Kernel slope was used as a summary measure of temporal weighting, with positive values indicating recency effects, negative values indicating primacy effects, and values near zero indicating approximately uniform integration across time.

## Supplementary information

This article contains supplementary information in Extended Data document.

## Acknowledgements

This work was funded by MICI-U/AEI/10.13039/501100011033, grant PID2023-147627OB-I00, and through the Severo Ochoa and María de Maeztu Program for Centers and Units of Excellence in R&D (CEX2020-001084-M). We thank CERCA Programme/Generalitat de Catalunya for institutional support.

## Declarations

The authors declare no competing interests.

## Appendix A

### Extended Data

**Fig. A1.**
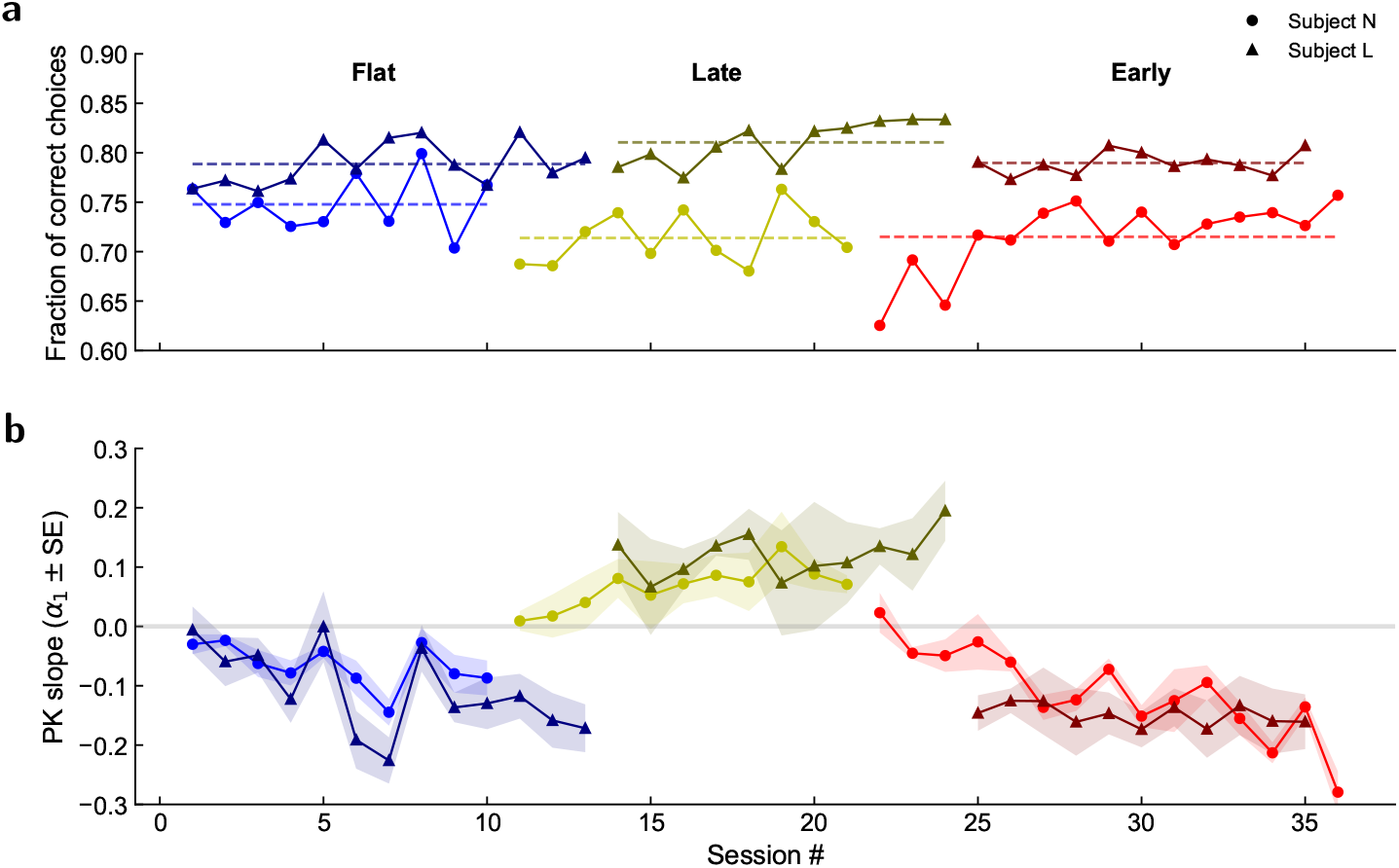
Choice behavior of the two subjects. **a**. Average fraction of choices reporting correct motion direction across sessions. **b**. Slope of the psychophysical kernels (±SE) for the seven pulses[14] shown in Fig. 1c, computed by regressing **y** = [*β*_1_, *β*_2_, …, *β*_7_] via **y** = *α*_0_ + *α*_1_**x, x** = [1, 2, 3, 4, 5, 6, 7].

**Fig. A2.**
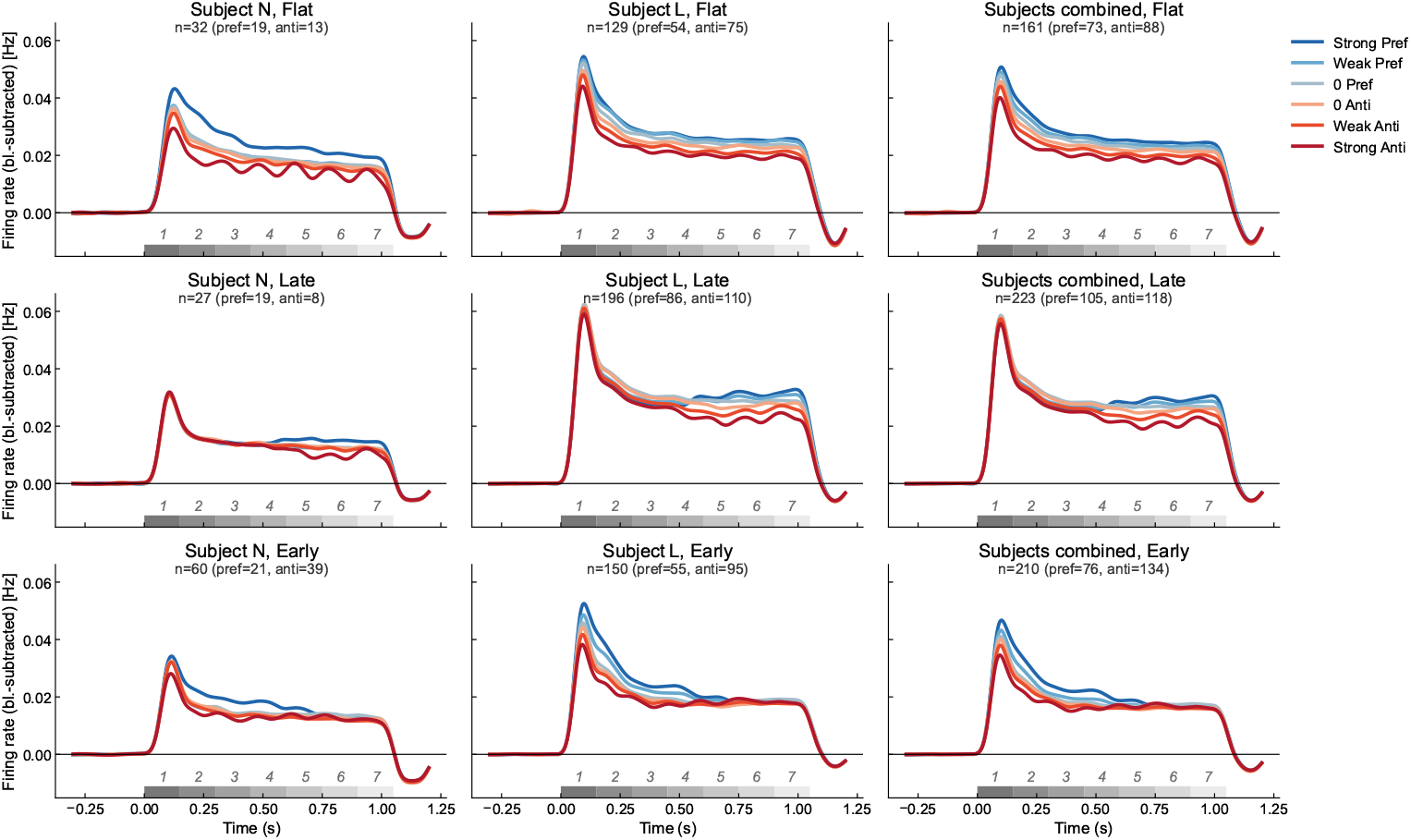
Showing results in 2a for the two subjects separately.

**Table A1.** Recording session dates grouped by subject and condition.

| Subject N |  |  | Subject L |  |  |
| --- | --- | --- | --- | --- | --- |
| Flat | Late | Early | Flat | Late | Early |
| 2016-07-26 | 2016-09-09 | 2016-10-19 | 2019-08-06 | 2019-10-08 | 2019-12-05 |
| 2016-08-02 | 2016-09-13 | 2016-10-20 | 2019-08-07 | 2019-10-09 | 2019-12-06 |
| 2016-08-03 | 2016-09-15 | 2016-10-21 | 2019-08-14 | 2019-10-10 | 2019-12-07 |
| 2016-08-04 | 2016-09-20 | 2016-10-27 | 2019-08-15 | 2019-10-16 | 2019-12-09 |
| 2016-08-12 | 2016-09-21 | 2016-10-28 | 2019-08-16 | 2019-10-21 | 2019-12-10 |
| 2016-08-23 | 2016-09-23 | 2016-11-01 | 2019-08-27 | 2019-10-23 | 2019-12-11 |
| 2016-08-25 | 2016-09-28 | 2016-11-02 | 2019-08-29 | 2019-10-25 | 2019-12-12 |
| 2016-08-31 | 2016-10-05 | 2016-11-03 | 2019-08-30 | 2019-11-20 | 2019-12-13 |
| 2016-09-06 | 2016-10-13 | 2016-11-09 | 2019-09-04 | 2019-11-22 | 2019-12-16 |
| 2016-09-08 | 2016-10-14 | 2016-11-10 | 2019-09-05 | 2019-11-26 | 2019-12-17 |
|  | 2016-10-18 | 2016-11-11 | 2019-09-16 | 2019-11-29 | 2019-12-18 |
|  |  | 2016-11-15 | 2019-09-17 |  |  |
|  |  | 2016-11-30 | 2019-09-19 |  |  |
|  |  | 2016-12-01 |  |  |  |
|  |  | 2016-12-02 |  |  |  |

**Table A2.** Trials per session and condition, correct trials and accuracy (%) summary.

| Subject | Condition | Sessions | Trials | Mean ± SEM | Correct | % ± SEM |
| --- | --- | --- | --- | --- | --- | --- |
| N | Flat | 10 | 13110 | 1311.0 ± 134.9 | 9809 | 74.8 ± 0.9 |
| N | Late | 11 | 16043 | 1458.5 ± 78.1 | 11465 | 71.4 ± 0.8 |
| N | Early | 15 | 17607 | 1173.8 ± 80.8 | 12541 | 71.5 ± 1.0 |
| L | Flat | 13 | 15410 | 1185.4 ± 37.1 | 12172 | 78.9 ± 0.6 |
| L | Late | 11 | 13173 | 1197.5 ± 21.4 | 10668 | 81.0 ± 0.7 |
| L | Early | 11 | 12469 | 1133.5 ± 50.8 | 9851 | 79.0 ± 0.4 |

**Table A3.** Trial distribution across coherence levels. Values are pooled across all sessions for each subject. The final column reports the percentage of trials at each coherence level and the session-wise SEM of that percentage.

| Subject | Avg. Coherence | Trials | Total | % ± SEM |
| --- | --- | --- | --- | --- |
| N | -50% | 6942 | 46760 | 14.85 ± 1.20 |
| N | -10% | 6453 | 46760 | 13.80 ± 1.80 |
| N | 0% | 19931 | 46760 | 42.63 ± 0.60 |
| N | +10% | 6510 | 46760 | 13.92 ± 1.20 |
| N | +50% | 6924 | 46760 | 14.81 ± 0.80 |
| L | -50% | 7627 | 41052 | 18.58 ± 0.60 |
| L | -10% | 7114 | 41052 | 17.33 ± 1.00 |
| L | 0% | 11315 | 41052 | 27.56 ± 0.80 |
| L | +10% | 7266 | 41052 | 17.70 ± 1.20 |
| L | +50% | 7730 | 41052 | 18.83 ± 0.90 |

**Table A4.** Psychometric regression parameters and *p*-values across conditions and subjects.

| Condition | Subject | $\beta_0 \pm$ SE | $p_{\beta_0}$ | $\beta_1 \pm$ SE | $p_{\beta_1}$ |
| --- | --- | --- | --- | --- | --- |
| Flat | N | +0.064 ± 0.019 | $8.70 \times 10^{-4}$ | 0.080 ± 0.002 | $< 10^{-300}$ |
| Flat | L | -0.475 ± 0.021 | $1.19 \times 10^{-115}$ | 0.051 ± 0.001 | $< 10^{-300}$ |
| Flat | Combined | -0.176 ± 0.014 | $2.03 \times 10^{-38}$ | 0.057 ± 0.001 | $< 10^{-300}$ |
| Late | N | +0.241 ± 0.017 | $1.52 \times 10^{-43}$ | 0.085 ± 0.002 | $< 10^{-300}$ |
| Late | L | -0.464 ± 0.023 | $7.03 \times 10^{-91}$ | 0.089 ± 0.002 | $< 10^{-300}$ |
| Late | Combined | -0.000 ± 0.013 | 0.994 | 0.084 ± 0.001 | $< 10^{-300}$ |
| Early | N | +0.034 ± 0.016 | 0.037 | 0.081 ± 0.002 | $< 10^{-300}$ |
| Early | L | -0.361 ± 0.022 | $3.61 \times 10^{-60}$ | 0.079 ± 0.002 | $< 10^{-300}$ |
| Early | Combined | -0.091 ± 0.013 | $2.25 \times 10^{-12}$ | 0.079 ± 0.001 | $< 10^{-300}$ |

**Table A5.** Neuron counts by subject and temporal-weighting condition splitting *d*′ ≥0.2 or *d*′ *<* 0.2, with *d*′ computed over the average firing rate during the whole stimulus presentation time. Numerical values indicate counts of units, with percentages of respective available units shown in parentheses.

| Subject |  | Flat | Late | Early | Total |
| --- | --- | --- | --- | --- | --- |
| N | $d' \geq 0.2$ | 21/32 (65.63%) | 10/27 (37.04%) | 25/60 (41.67%) | 56/119 (47.06%) |
| L | $d' \geq 0.2$ | 76/129 (58.91%) | 104/196 (53.06%) | 117/150 (78.00%) | 297/475 (62.53%) |
| N | $d' < 0.2$ | 11/32 (34.38%) | 17/27 (62.96%) | 35/60 (58.33%) | 63/119 (52.94%) |
| L | $d' < 0.2$ | 53/129 (41.09%) | 92/196 (46.94%) | 33/150 (22.00%) | 178/475 (37.47%) |

**Fig. A3.**
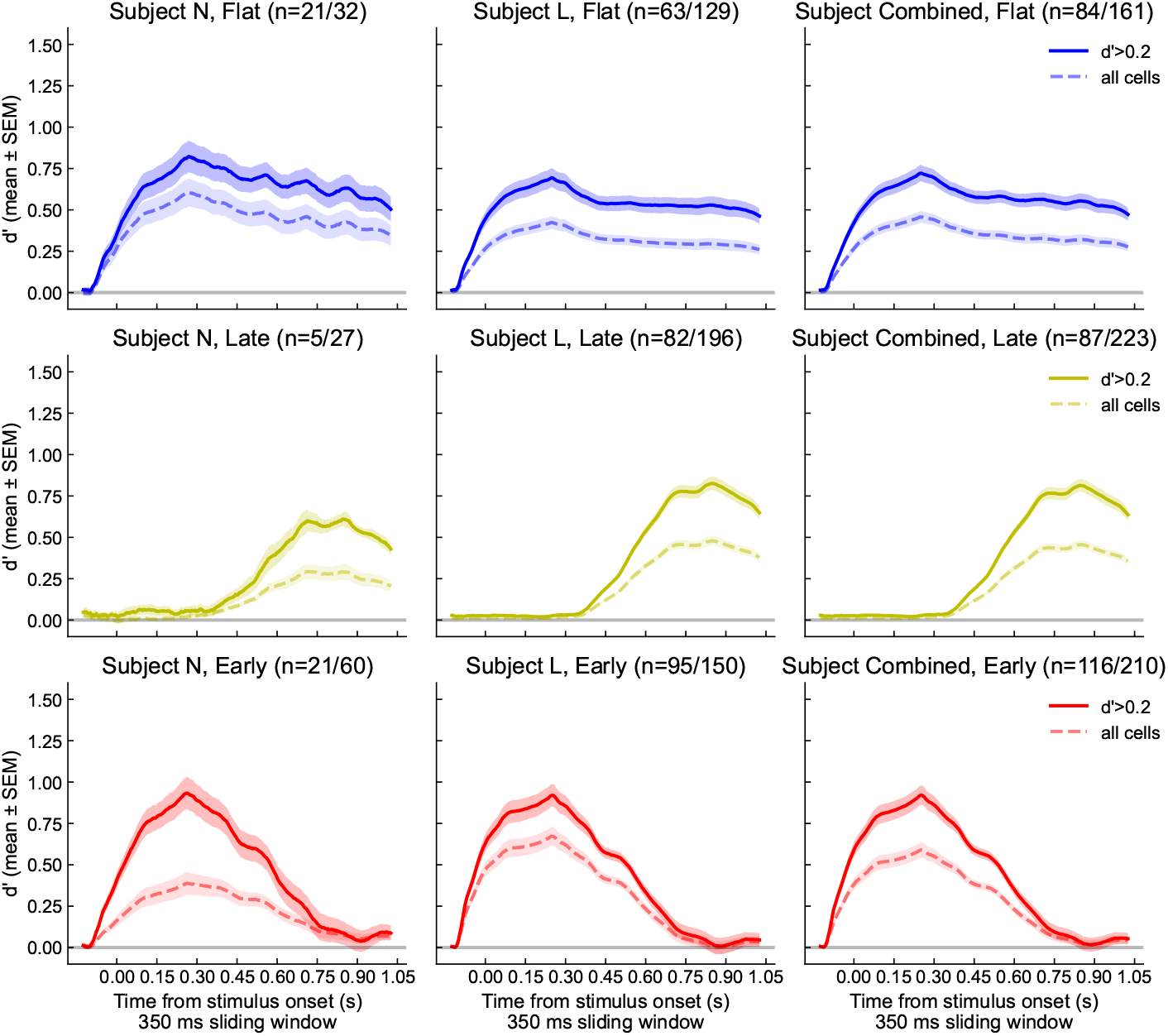
Subject-specific d′. Showing results in 2b for the two subjects separately. In parentheses, we indicate the number of cells showing epoch-wise *d*′ *>* 0.2 out of the total number of cells available.

**Fig. A4.**
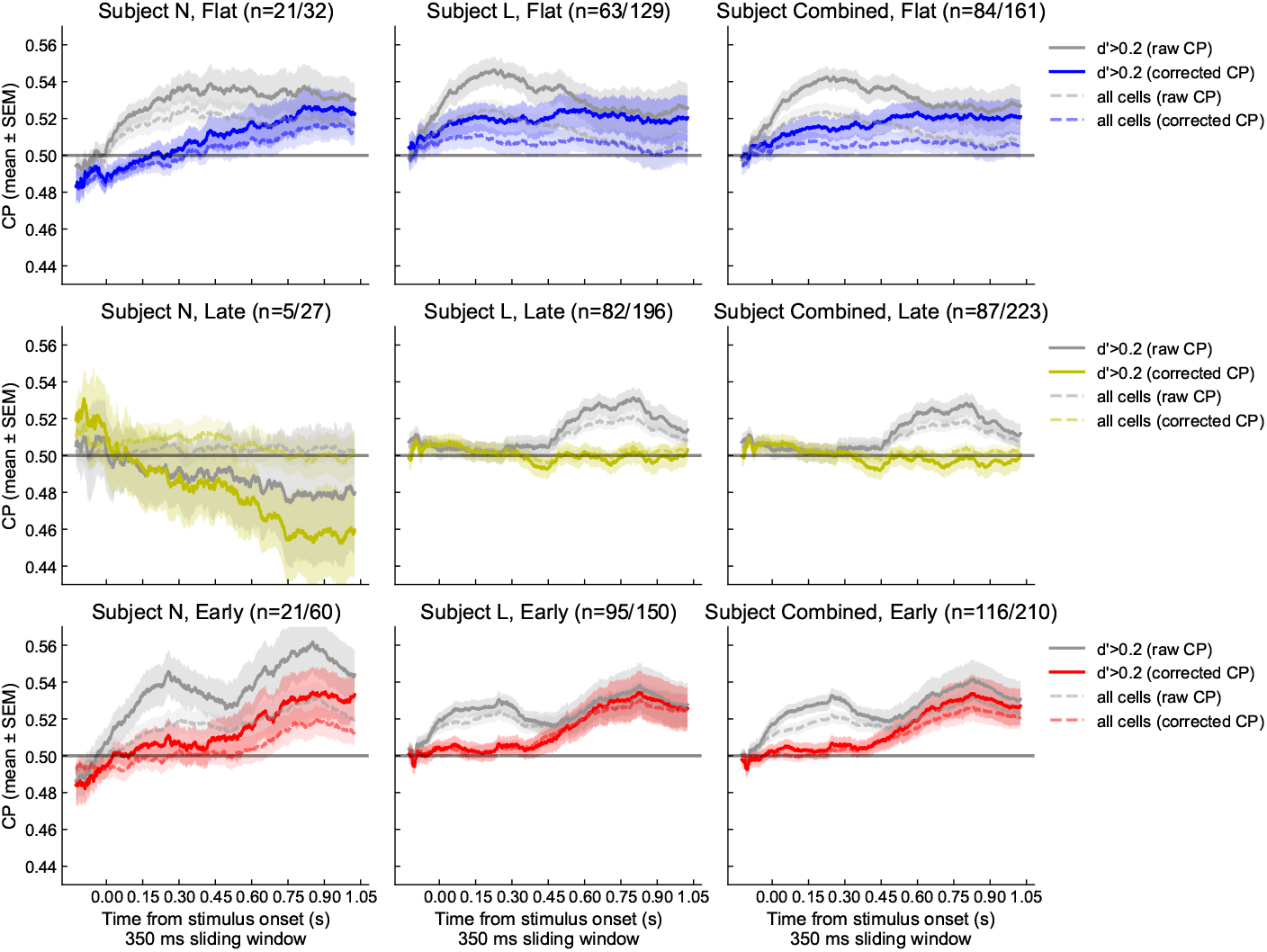
Subject-specific CP. Showing results in 2c for all available cells and for the two subjects separately. In parentheses, cells showing epoch-wise *d*′ *>* 0.2 out of the total number of cells available.

**Fig. A5.**
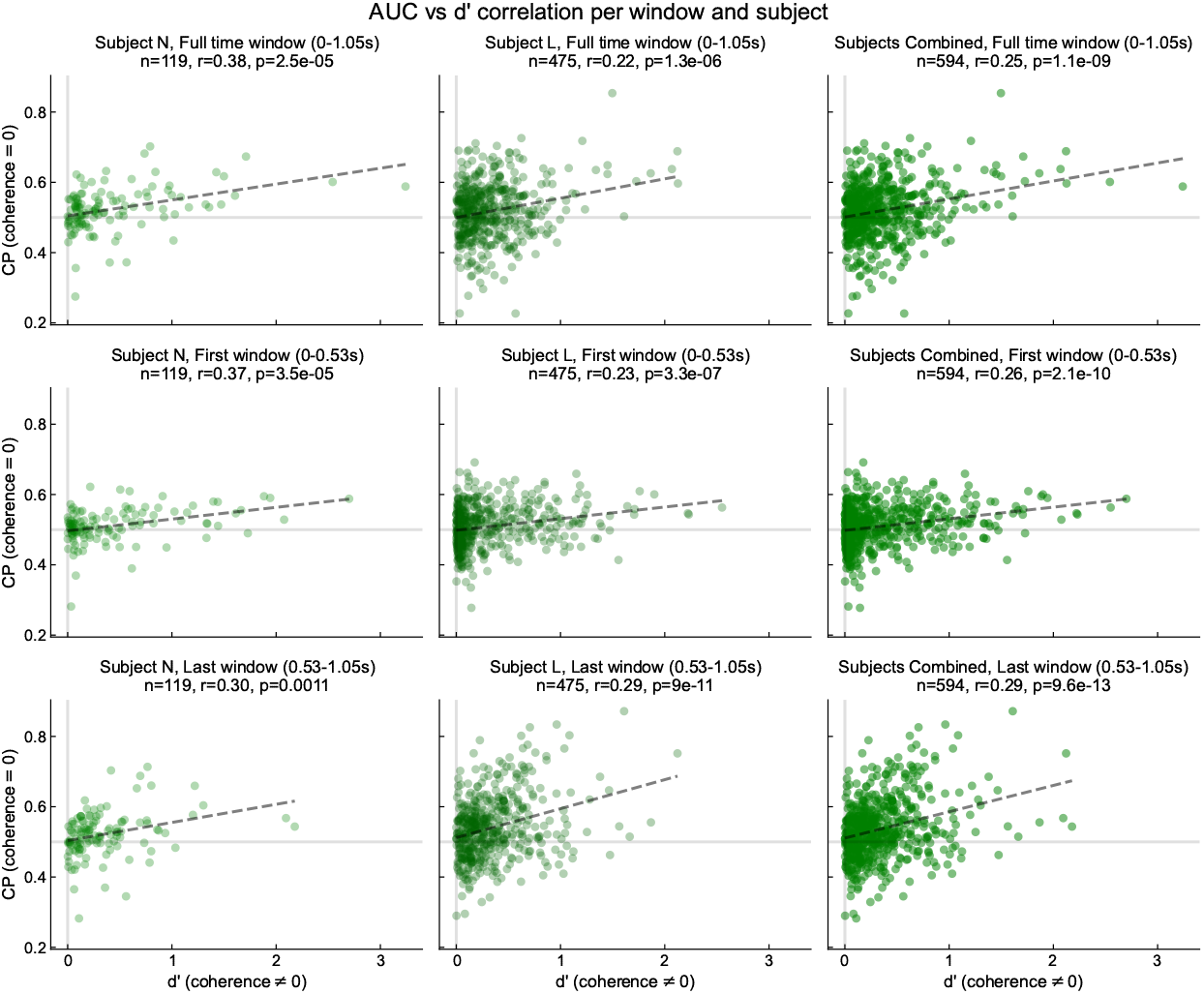
Correlation between CP raw and d′.

**Fig. A6.**
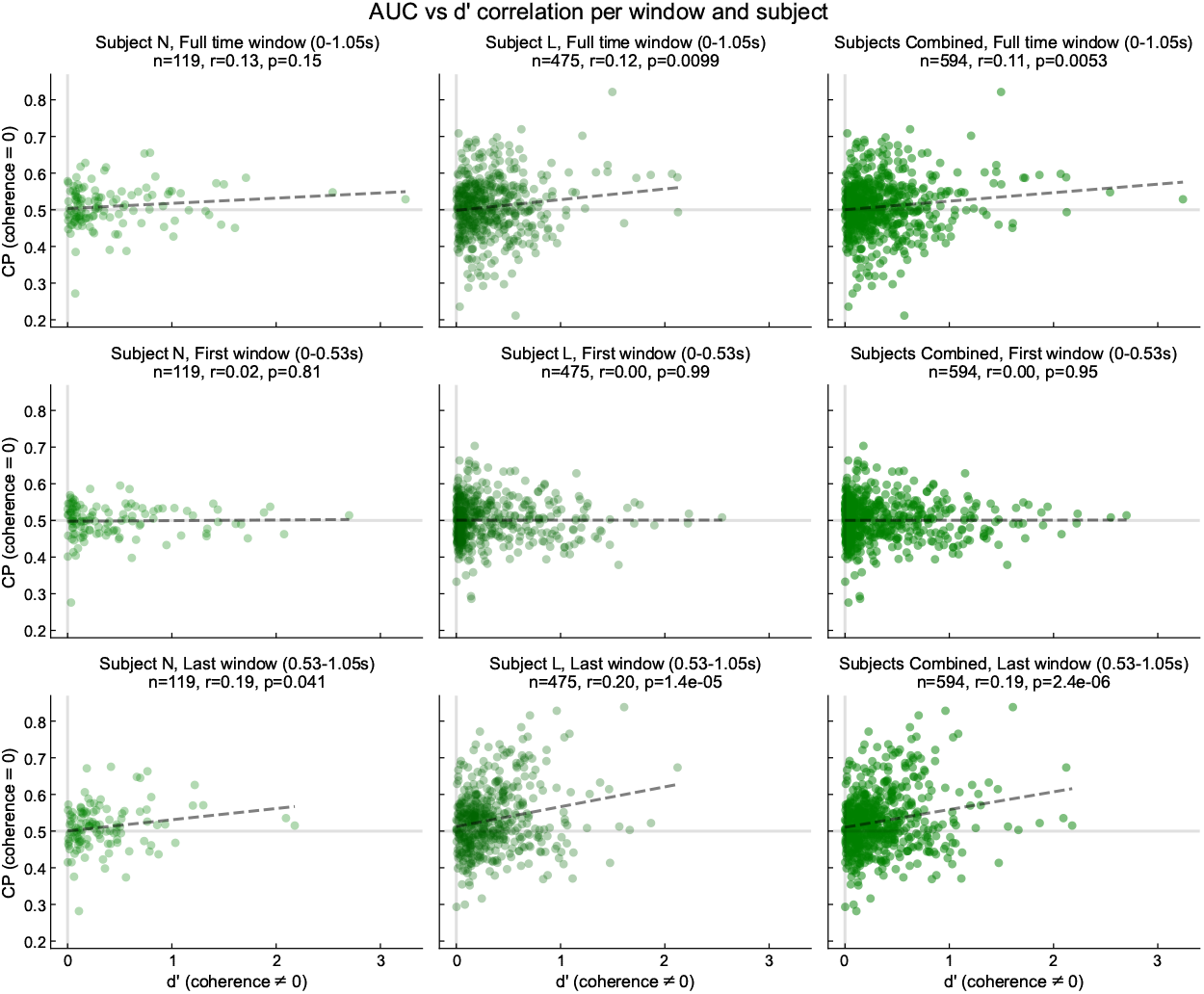
Correlation between CP corrected and d′.

**Fig. A7.**
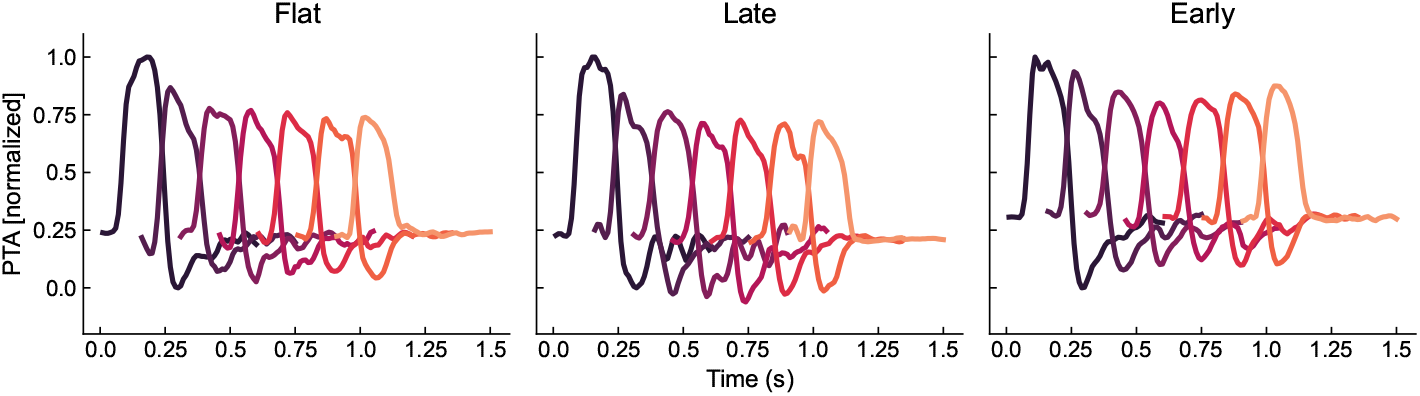
PTA evoked by motion pulses in MT data. PTA averaged across sessions for each task condition, computed over the weighted sum of firing rates for cells recorded in each session.

**Fig. A8.**
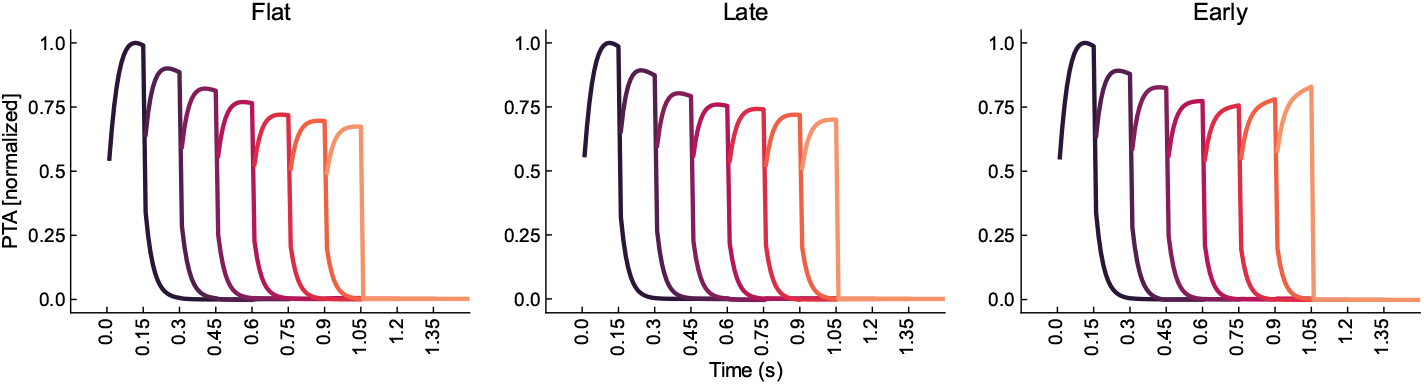
PTA evoked by motion pulses in the neural network model. PTA averaged across sessions for each task condition, computed over the weighted sum of firing rate simulated.

**Fig. A9.**
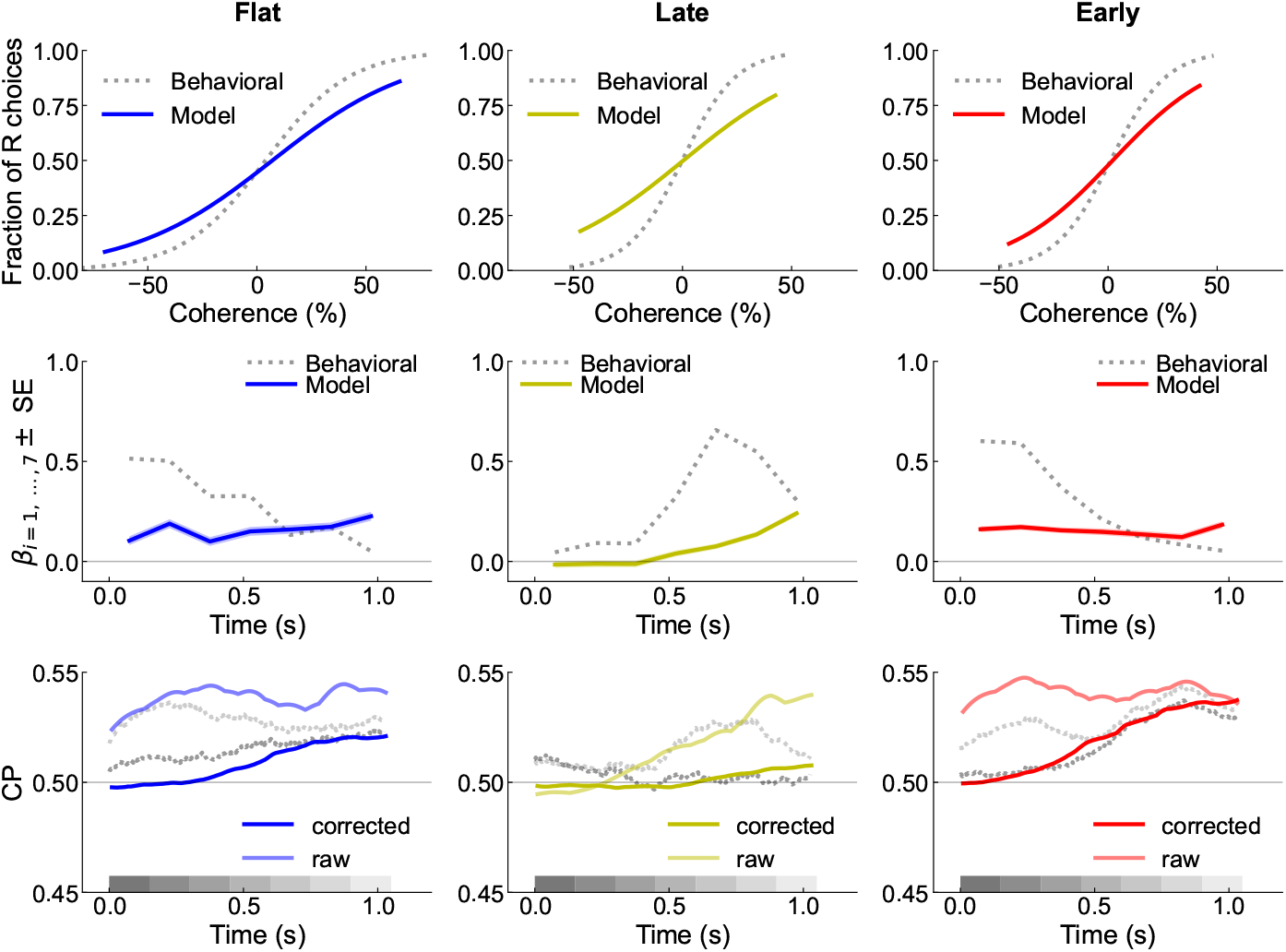
Equivalent results for weaker feed-forward conntectivity. Same as Fig. 3 but tuning connectivity weights to get raw CP (shaded colors in bottom row) within ranges comparable with MT data (gray in bottom row). We set *J*_FF,1_ = *J*_FF,2_ = 0.15 to show that it drastically impacts choice estimation (top row) and the magnitude of PKs (middle row). We also set the *J*_FB,1_ = *J*_FB,2_ = 0.001 to match such changes and recover the qualitative shape of raw and corrected CPs.

**Fig. A10.**
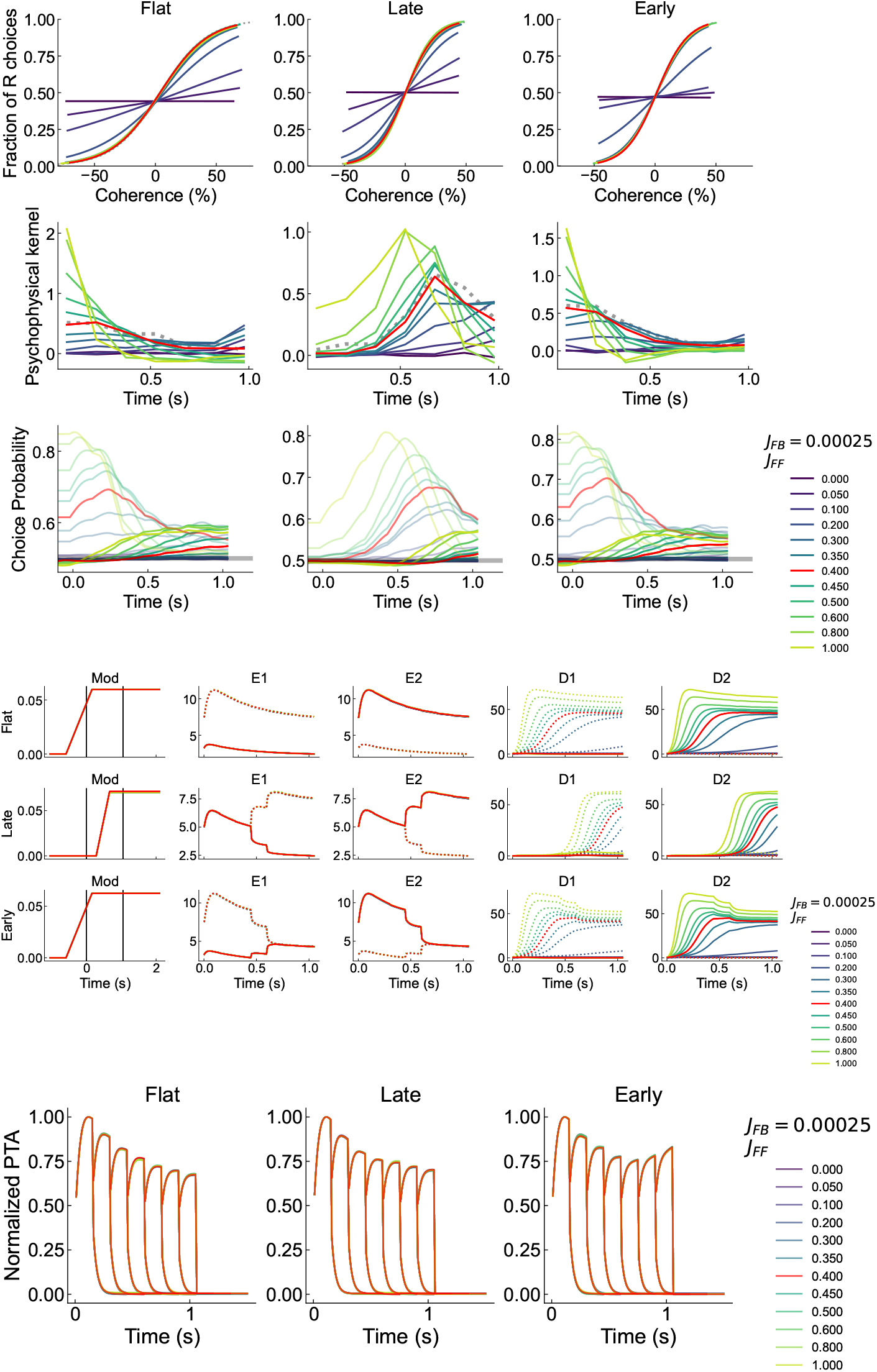
Effects of varying *J*_*FF*_. Red indicates model output for optimized parameters. In CP plots, dotted lines indicate raw CP, solid lines indicate corrected CP. In firing rate plots, dotted lines indicate tuning to target 1, solid lines indicate tuning to target 2.

**Fig. A11.**
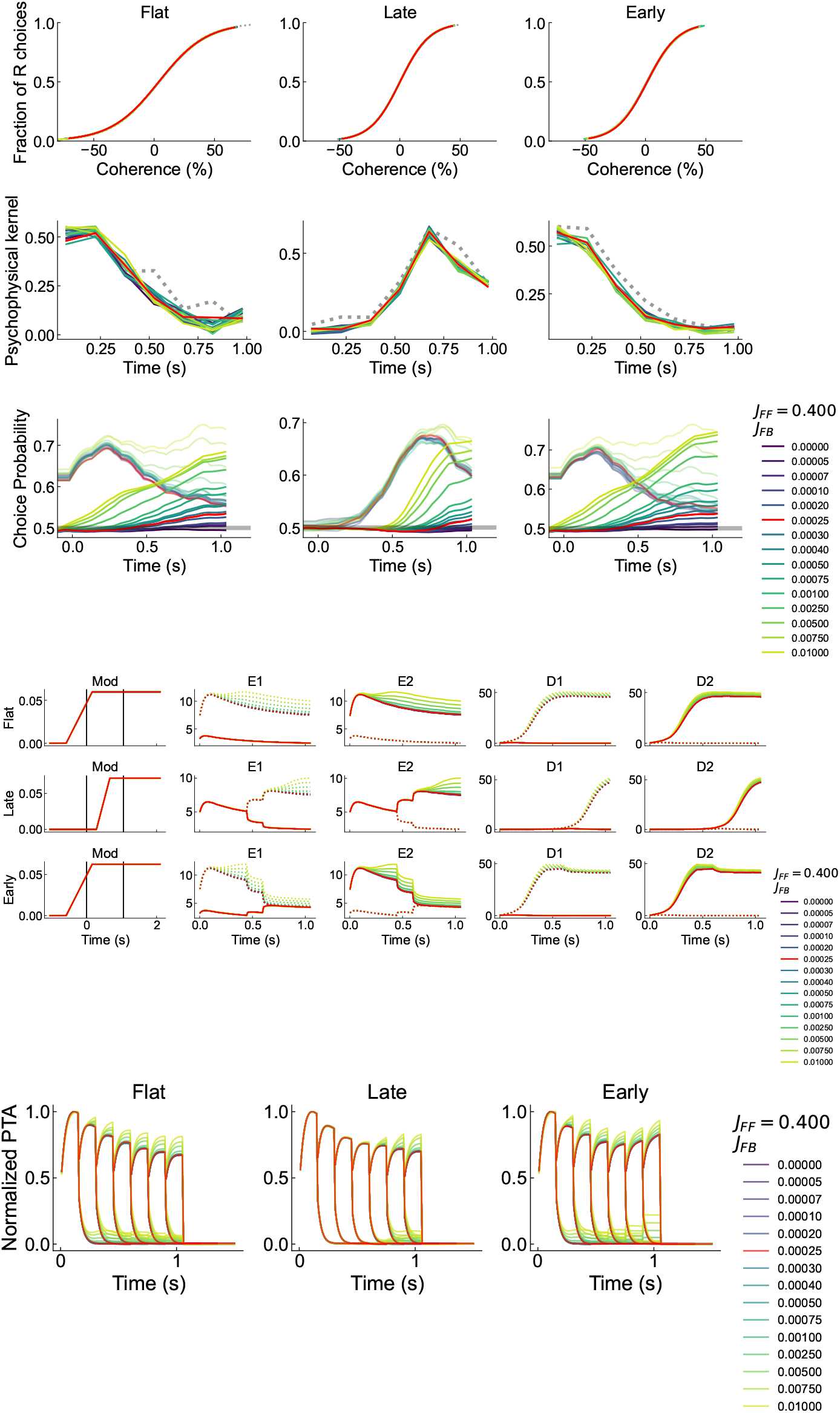
Effects of varying *J*_*FB*_. Red indicates model output for optimized parameters. In CP plots, dotted lines indicate raw CP, solid lines indicate corrected CP. In firing rate plots, dotted lines indicate tuning to target 1, solid lines indicate tuning to target 2.

**Fig. A12.**
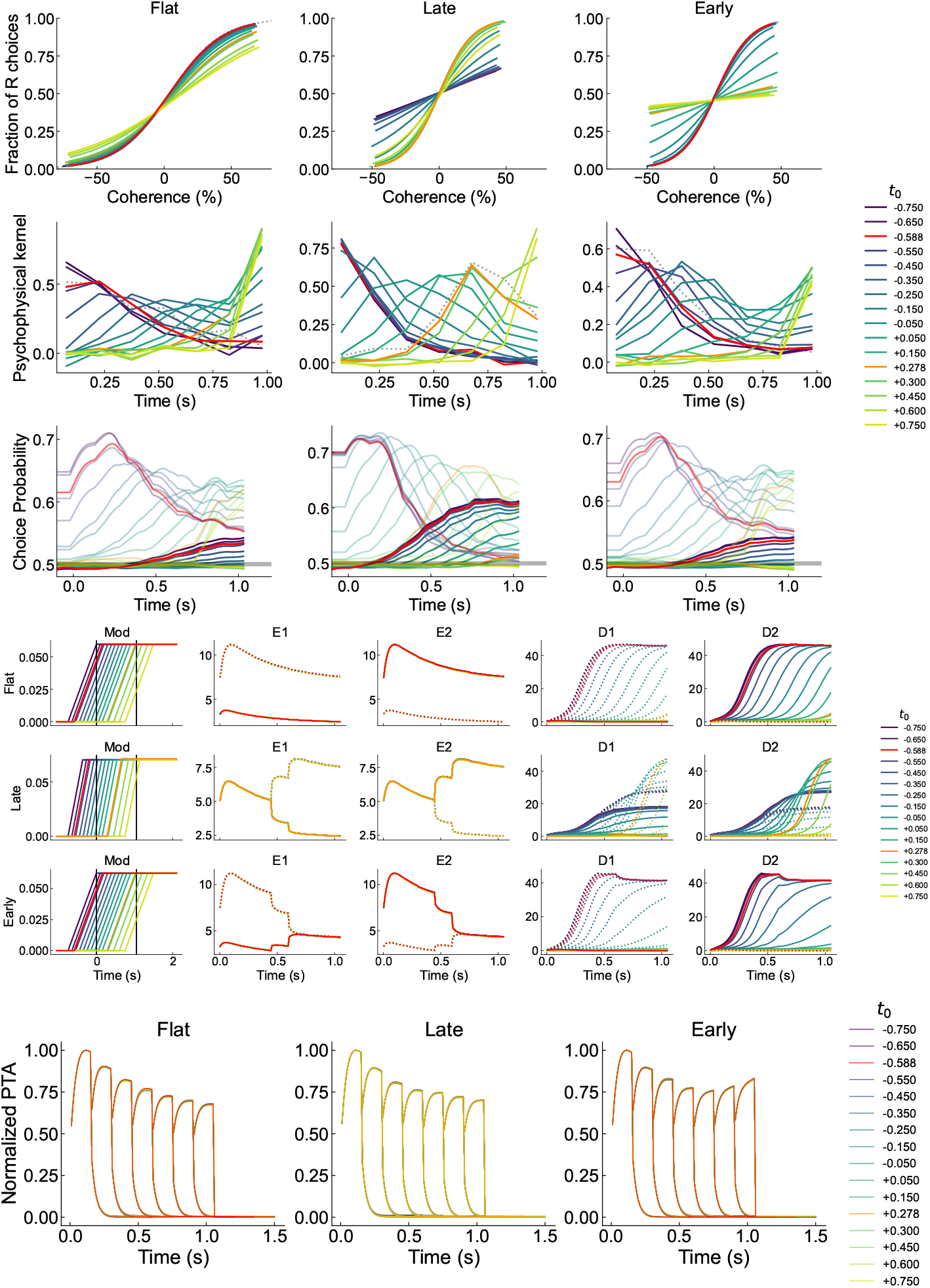
Effects of varying *t*_0,*e*_ + *t*_offset_, *t*_offset_ ∈ − [1.05, +1.15] (s) in *e* = {flat, late, early}. Red indicates model output for optimized parameter in flat and early conditions; orange indicates model output for optimized parameter in late condition. In CP plots, dotted lines indicate raw CP, solid lines indicate corrected CP. In firing rate plots, dotted lines indicate tuning to target 1, solid lines indicate tuning to target 2.

**Fig. A13.**
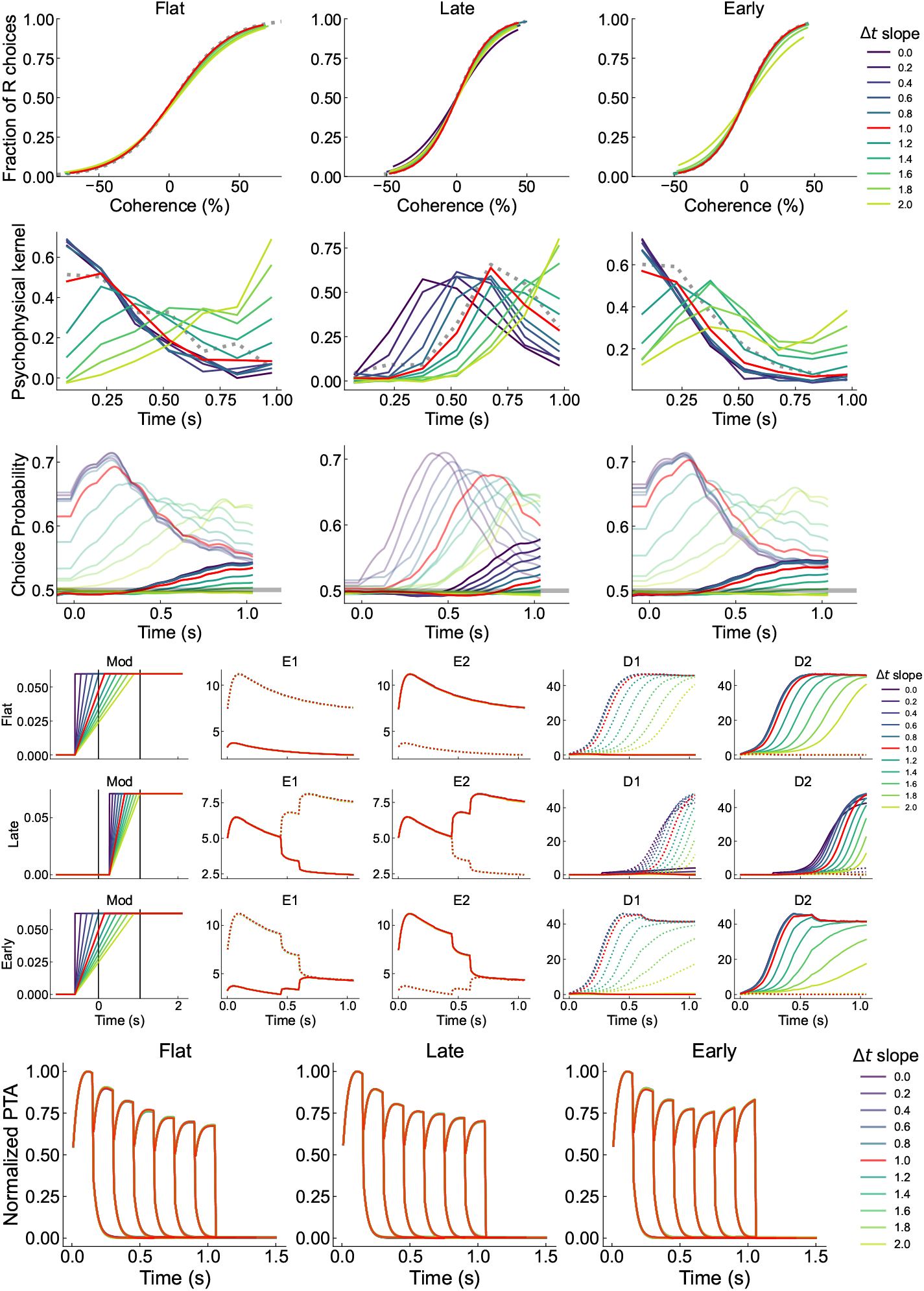
Effects of varying *I*_mod_ slope as *k*Δ*t*_*e*_, *k* ∈ [0, 2] (s) for *e* = {flat, late, early}. Red indicates model output for optimized Δ*t*_*e*_ (*k* = 1). In CP plots, dotted lines indicate raw CP, solid lines indicate corrected CP. In firing rate plots, dotted lines indicate tuning to target 1, solid lines indicate tuning to target 2.

**Fig. A14.**
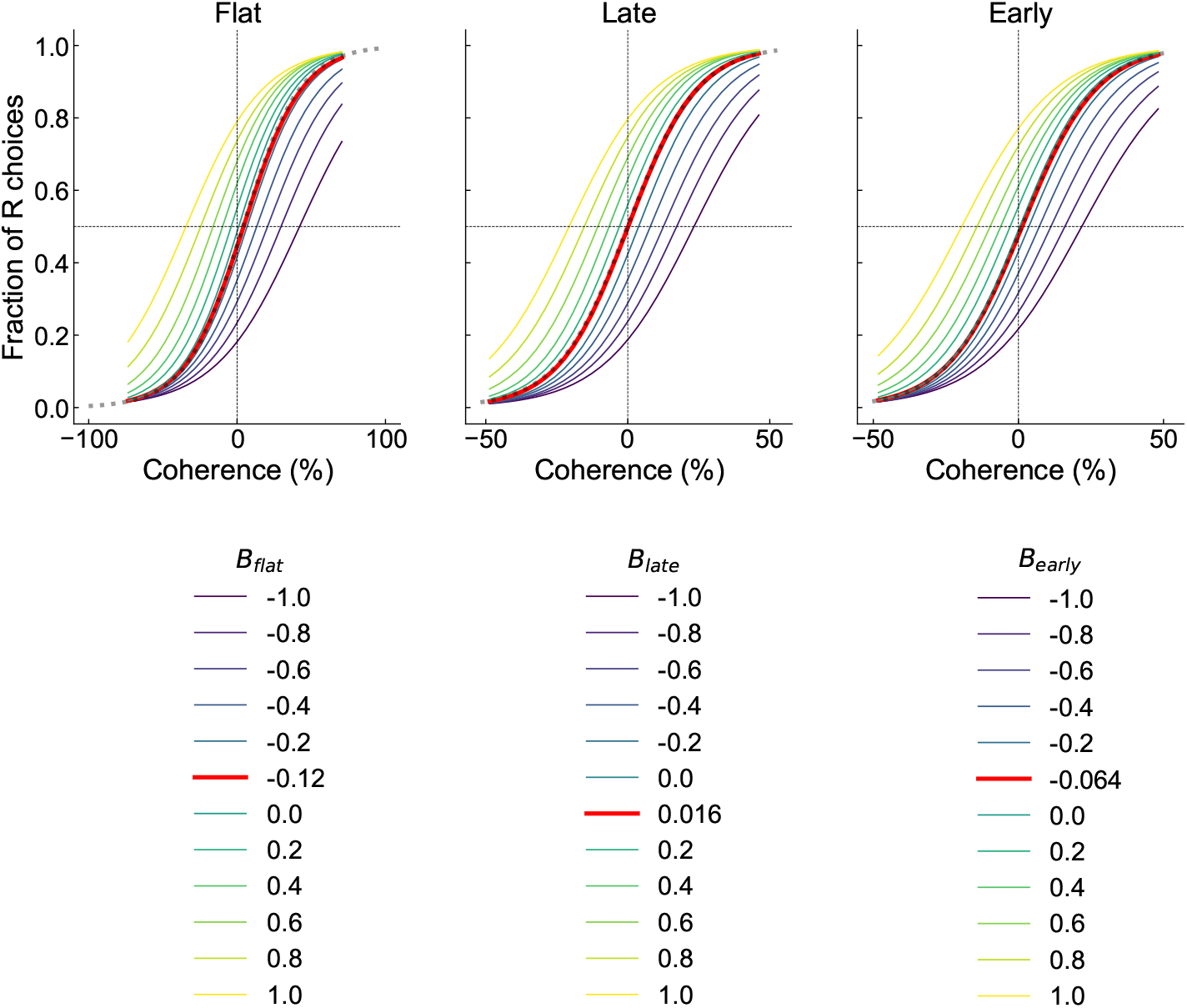
Effects of varying *B*_flat_, *B*_late_, *B*_early_ on choice estimation. The parameters and *σ*_flat_ = 0.498, *σ*_late_ = 0.531, *σ*_early_ = 0.583 were kept fixed in the respective condition. Red indicates model output for optimized parameters; dotted lines indicate logistic fits on behavioral data.

**Fig. A15.**
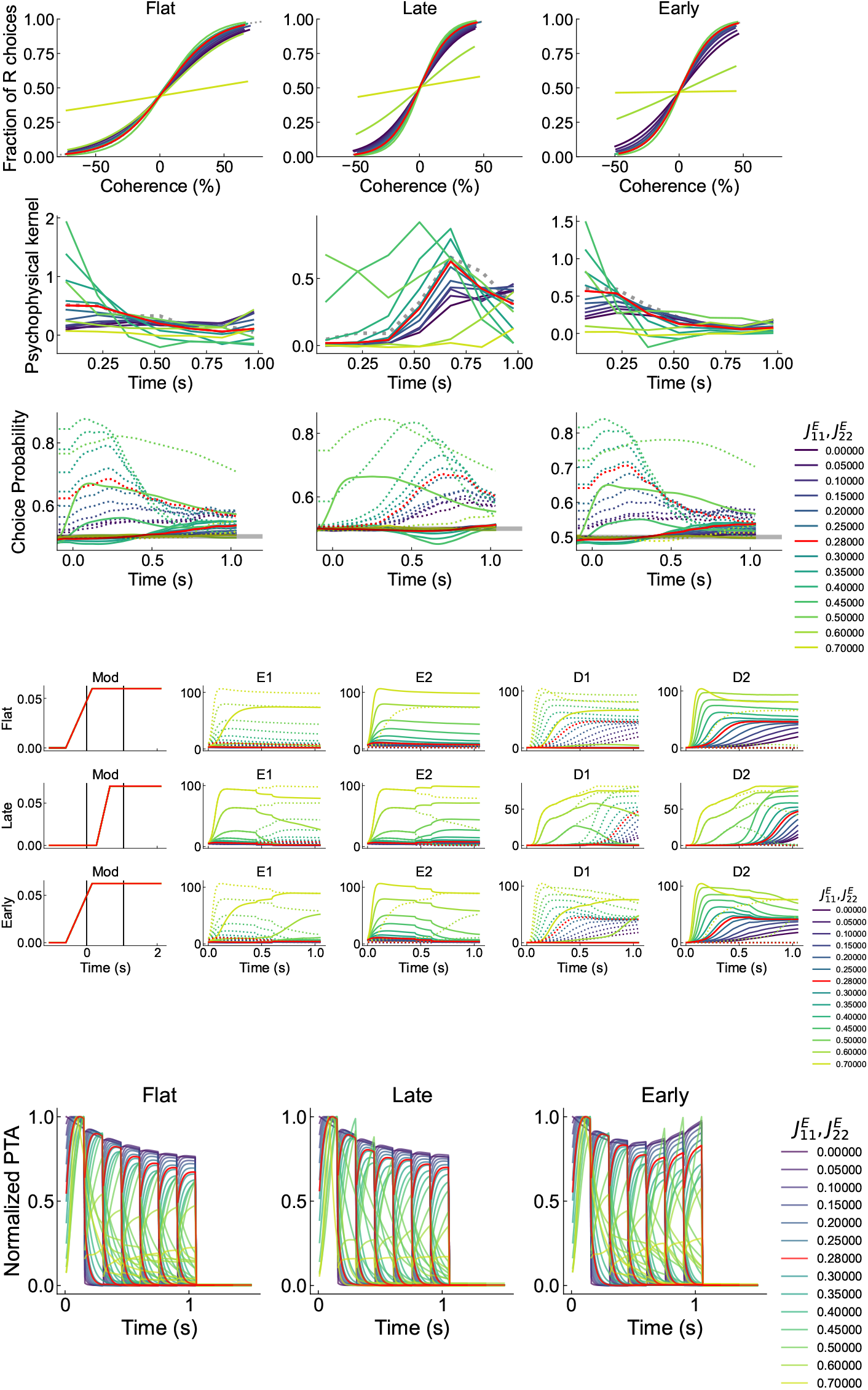
Effects of varying 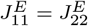. Red indicates model output for optimized parameters. In CP plots, dotted lines indicate raw CP, solid lines indicate corrected CP. In firing rate plots, dotted lines indicate tuning to target 1, solid lines indicate tuning to target 2.

**Fig. A16.**
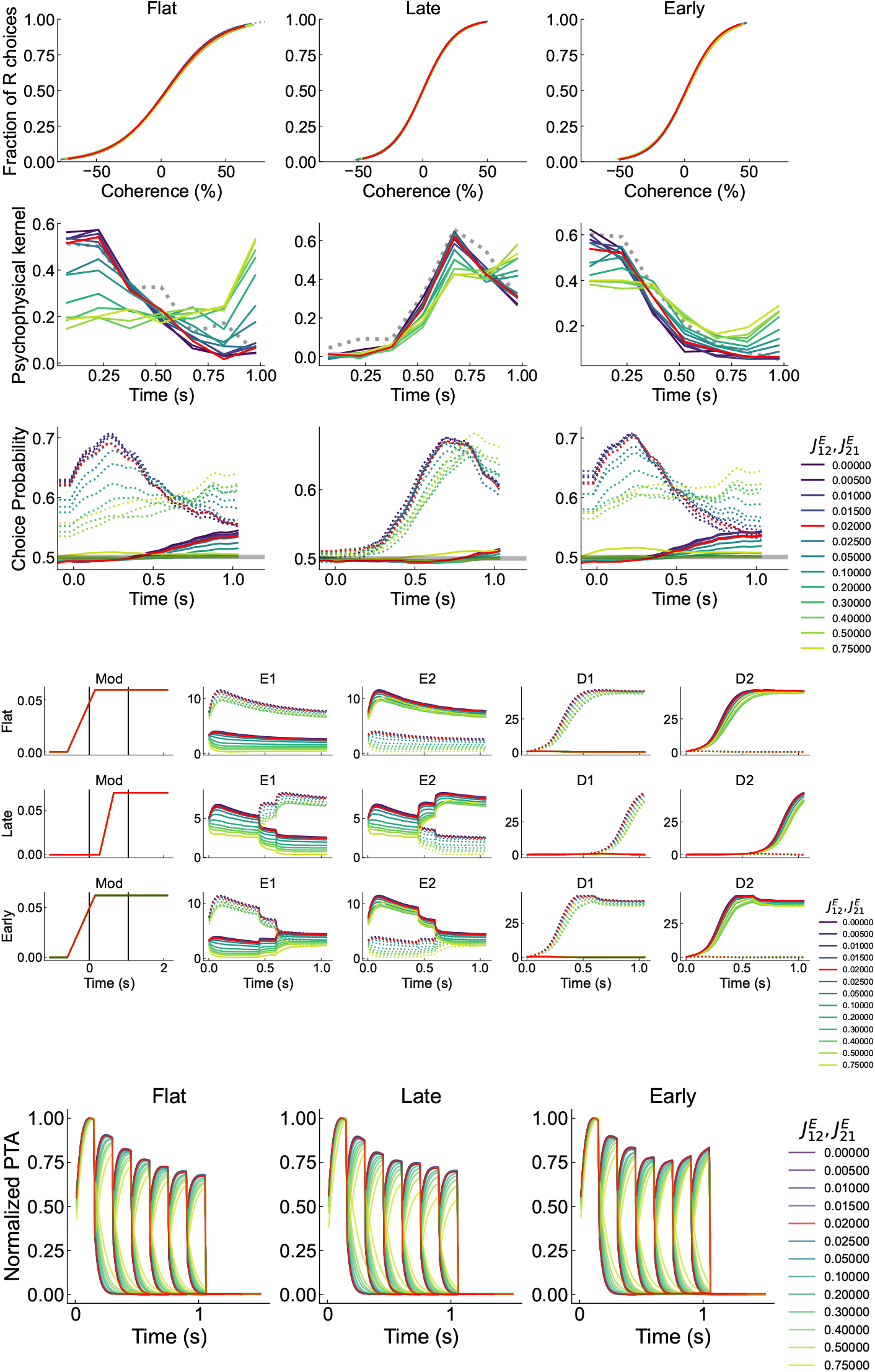
Effects of varying 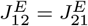. Red indicates model output for optimized parameters. In CP plots, dotted lines indicate raw CP, solid lines indicate corrected CP. In firing rate plots, dotted lines indicate tuning to target 1, solid lines indicate tuning to target 2.

**Fig. A17.**
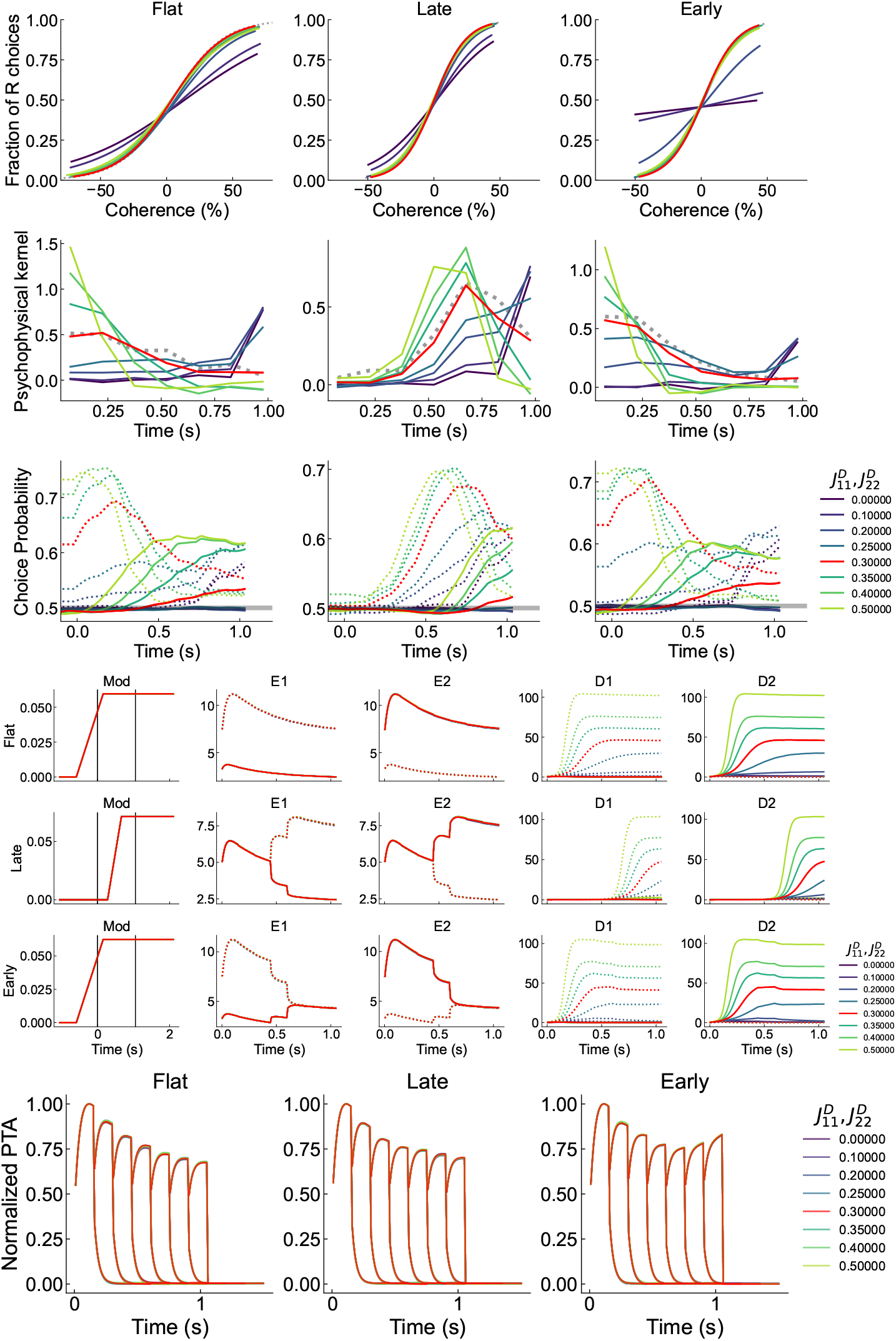
Effects of varying 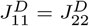. Red indicates model output for optimized parameters. In CP plots, dotted lines indicate raw CP, solid lines indicate corrected CP. In firing rate plots, dotted lines indicate tuning to target 1, solid lines indicate tuning to target 2.

**Fig. A18.**
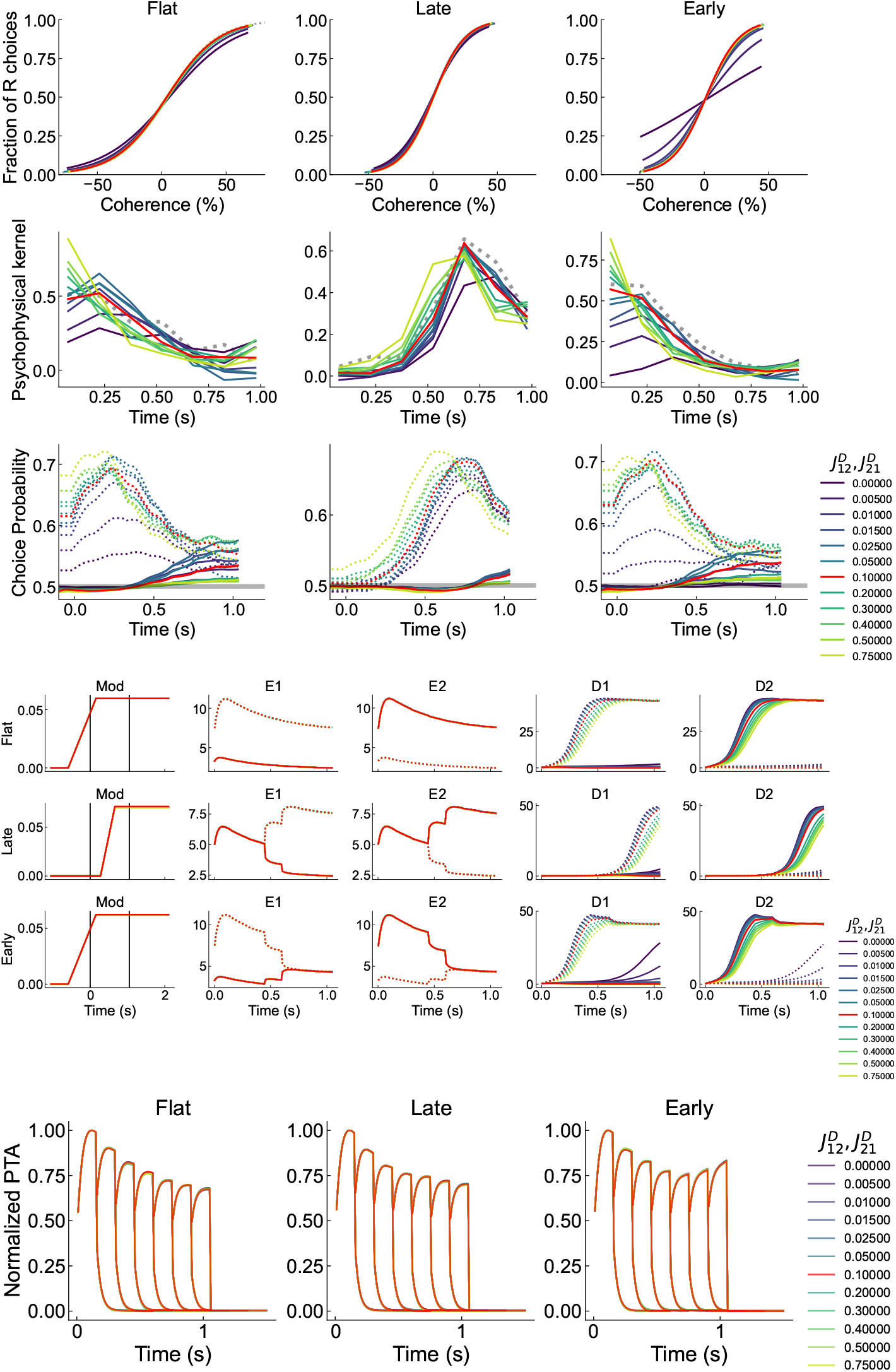
Effects of varying 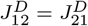. Red indicates model output for optimized parameters. In CP plots, dotted lines indicate raw CP, solid lines indicate corrected CP. In firing rate plots, dotted lines indicate tuning to target 1, solid lines indicate tuning to target 2.

**Fig. A19.**
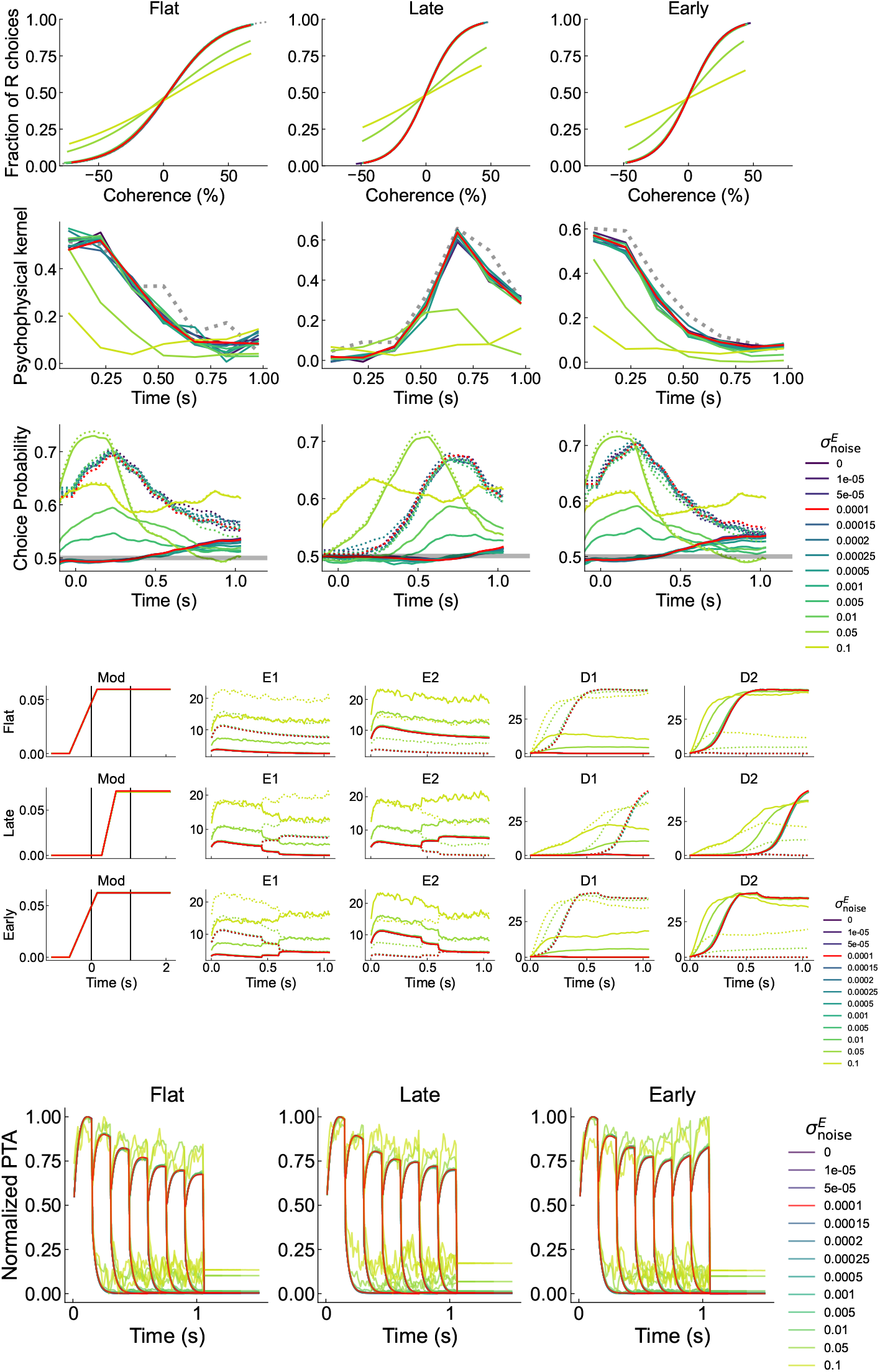
Effects of varying 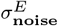. Red indicates model output for optimized parameters. In CP plots, dotted lines indicate raw CP, solid lines indicate corrected CP. In firing rate plots, dotted lines indicate tuning to target 1, solid lines indicate tuning to target 2.

**Fig. A20.**
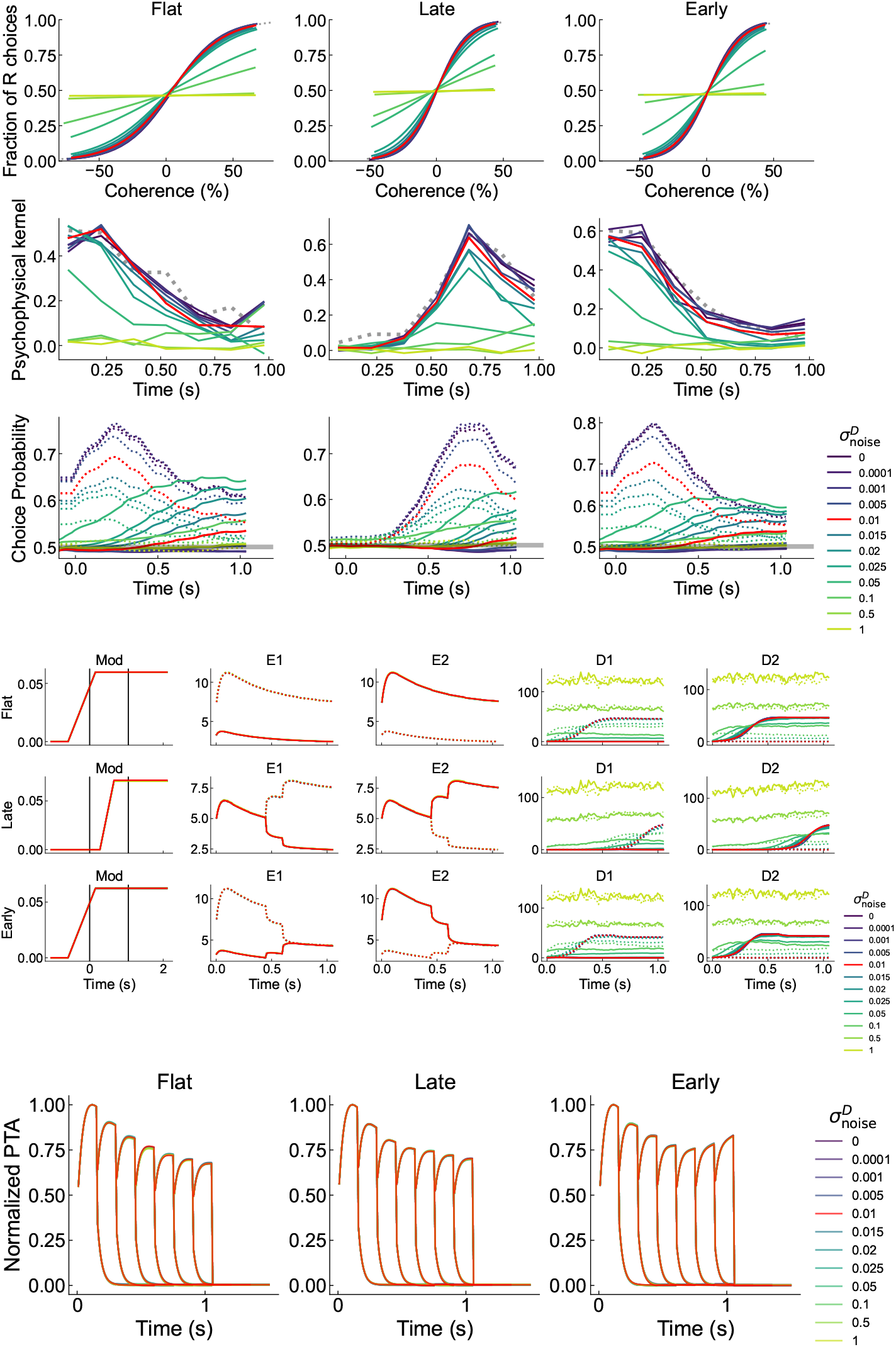
Effects of varying 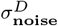. Red indicates model output for optimized parameters. In CP plots, dotted lines indicate raw CP, solid lines indicate corrected CP. In firing rate plots, dotted lines indicate tuning to target 1, solid lines indicate tuning to target 2.

## References

[1] Albright, T.D.: Direction and orientation selectivity of neurons in visual area mt of the macaque. Journal of neurophysiology 52(6), 1106–1130 (1984)

[2] Allman, J., Miezin, F., McGuinness, E.: Direction-and velocity-specific responses from beyond the classical receptive field in the middle temporal visual area (mt). Perception 14(2), 105–126 (1985)

[3] Britten, K.H., Shadlen, M.N., Newsome, W.T., Movshon, J.A.: The analysis of visual motion: a comparison of neuronal and psychophysical performance. Journal of Neuroscience 12(12), 4745–4765 (1992)

[4] Shadlen, M.N., Newsome, W.T.: Neural basis of a perceptual decision in the parietal cortex (area lip) of the rhesus monkey. Journal of neurophysiology 86(4), 1916–1936 (2001)

[5] Rorie, A.E., Gao, J., McClelland, J.L., Newsome, W.T.: Integration of sensory and reward information during perceptual decision-making in lateral intraparietal cortex (lip) of the macaque monkey. PloS one 5(2), 9308 (2010)

[6] Roitman, J.D., Shadlen, M.N.: Response of neurons in the lateral intraparietal area during a combined visual discrimination reaction time task. Journal of Neuroscience 22(21), 9475–9489 (2002)

[7] Steinemann, N., Stine, G.M., Trautmann, E., Zylberberg, A., Wolpert, D.M., Shadlen, M.N.: Direct observation of the neural computations underlying a single decision. eLife 12 (2024) 10.7554/eLife.90859

[8] Ferrera, V.P., Yanike, M., Cassanello, C.: Frontal eye field neurons signal changes in decision criteria. Nature neuroscience 12(11), 1458–1462 (2009)

[9] Costello, M.G., Zhu, D., Salinas, E., Stanford, T.R.: Perceptual modulation of motor—but not visual—responses in the frontal eye field during an urgent-decision task. Journal of neuroscience 33(41), 16394–16408 (2013)

[10] Coe, B., Tomihara, K., Matsuzawa, M., Hikosaka, O.: Visual and anticipatory bias in three cortical eye fields of the monkey during an adaptive decision-making task. Journal of Neuroscience 22(12), 5081–5090 (2002)

[11] Mazurek, M.E., Roitman, J.D., Ditterich, J., Shadlen, M.N.: A role for neural integrators in perceptual decision making. Cerebral cortex 13(11), 1257–1269 (2003)

[12] Yates, J.L., Park, I.M., Katz, L.N., Pillow, J.W., Huk, A.C.: Functional dissection of signal and noise in mt and lip during decision-making. Nature neuroscience 20(9), 1285–1292 (2017)

[13] Levi, A.J., Yates, J.L., Huk, A.C., Katz, L.N.: Strategic and dynamic temporal weighting for perceptual decisions in humans and macaques. eneuro 5(5) (2018)

[14] Levi, A.J., Zhao, Y., Park, I.M., Huk, A.C.: Sensory and choice responses in mt distinct from motion encoding. Journal of Neuroscience 43(12), 2090–2103 (2023)

[15] Gold, J.I., Shadlen, M.N.: The neural basis of decision making. Annual Review of Neuroscience 30, 535–574 (2007)

[16] Simoncelli, E.P., Heeger, D.J.: A model of neuronal responses in visual area mt. Vision Research 38(5), 743–761 (1998)

[17] Rust, N.C., Mante, V., Simoncelli, E.P., Movshon, J.A.: How mt cells analyze the motion of visual patterns. Nature Neuroscience 9(11), 1421–1431 (2006)

[18] Ratcliff, R., McKoon, G.: The diffusion decision model: theory and data for two-choice decision tasks. Neural Computation 20(4), 873–922 (2008)

[19] Bogacz, R., Brown, E., Moehlis, J., Holmes, P., Cohen, J.D.: The physics of optimal decision making: A formal comparison of models of performance in two-alternative forced-choice tasks. Psychological Review 113(4), 700–765 (2006)

[20] Krajbich, I., Rangel, A.: Multialternative drift-diffusion model predicts the relationship between visual fixations and choice in value-based decisions. Proceedings of the National Academy of Sciences 108(33), 13852–13857 (2011)

[21] Wang, X.-J.: Probabilistic decision making by slow reversion in cortical circuits. Neuron 36(5), 955–968 (2002)

[22] Wong, K.-F., Wang, X.-J.: A recurrent network mechanism of time integration in perceptual decisions. Journal of Neuroscience 26(4), 1314–1328 (2006)

[23] Prat-Ortega, G., Wimmer, K., Roxin, A., Rocha, J.: Flexible categorization in perceptual decision making. Nature Communications 12(1), 1283 (2021) 10.1038/s41467-021-21501-z

[24] Diederich, A., Busemeyer, J.R.: Modeling the effects of payoff on response bias in a perceptual discrimination task: Bound-change, drift-rate-change, or two-stage-processing hypothesis. Perception & Psychophysics 68(2), 194–207 (2006)

[25] Krajbich, I., Armel, C., Rangel, A.: Visual fixations and the computation and comparison of value in simple choice. Nature Neuroscience 13(10), 1292–1298 (2010)

[26] Purcell, B.A., Kiani, R.: Hierarchical decision processes that operate over distinct timescales underlie choice and changes in strategy. Proceedings of the national academy of sciences 113(31), 4531–4540 (2016)

[27] Law, C.T., Gold, J.I.: Neural correlates of perceptual learning in a sensory-motor, but not a sensory, cortical area. Nature Neuroscience 11(5), 505–513 (2008)

[28] Wimmer, K., Compte, A., Roxin, A., Peixoto, D., Renart, A., De La Rocha, J.: Sensory integration dynamics in a hierarchical network explains choice probabilities in cortical area mt. Nature communications 6(1), 6177 (2015)

[29] Britten, K.H., Newsome, W.T., Shadlen, M.N., Celebrini, S., Movshon, J.A.: A relationship between behavioral choice and the visual responses of neurons in macaque mt. Visual neuroscience 13(1), 87–100 (1996)

[30] Shadlen, M.N., Britten, K.H., Newsome, W.T., Movshon, J.A.: A computational analysis of the relationship between neuronal and behavioral responses to visual motion. The Journal of Neuroscience 16(4), 1486–1510 (1996) 10.1523/JNEUROSCI.16-04-01486.1996

[31] Haefner, R.M., Gerwinn, S., Macke, J.H., Bethge, M.: Inferring decoding strategies from choice probabilities in the presence of correlated variability. Nature Neuroscience 16(2), 235–242 (2013) 10.1038/nn.3309

[32] Nienborg, H., Cumming, B.G.: Decision-related activity in sensory neurons reflects more than a neuron’s causal effect. Nature 459(7243), 89–92 (2009)

[33] Pletenev, A., Born, R.T., DeAngelis, G.C., Doudlah, R., Fujita, I., Gold, J.I., Goris, R.L.T., Huk, A.C., Krug, K., Laamerad, P., al.: Reassessing choice probability: What 59 macaque studies tell us about decision-related activity in visual cortex. BioRxiv (2026) 10.64898/2026.06.08.730961

[34] Kiani, R., Hanks, T.D., Shadlen, M.N.: Bounded integration in parietal cortex underlies decisions even when viewing duration is dictated by the environment. The Journal of Neuroscience 28(12), 3017–3029 (2008) 10.1523/JNEUROSCI.4761-07.2008

[35] Roach, J.P., Churchland, A.K., Engel, T.A.: Choice selective inhibition drives stability and competition in decision circuits. Nature communications 14(1), 147 (2023)

[36] Quinn, K.R., Seillier, L., Butts, D.A., Nienborg, H.: Decision-related feedback in visual cortex lacks spatial selectivity. Nature Communications 12(1), 4473 (2021)

[37] Giarrocco, F., Averbeck, B.B.: Neuronal dynamics, timing, and flow of sensory and choice-related information in auditory-prefrontal circuitry. Nature Communications (2026)

[38] Ishizu, K., Nishimoto, S., Ueoka, Y., Funamizu, A.: Localized and global representation of prior value, sensory evidence, and choice in male mouse cerebral cortex. Nature Communications 15(1), 4071 (2024)

[39] Ferro, D., Cash-Padgett, T., Wang, M.Z., Hayden, B.Y., Moreno-Bote, R.: Gaze-centered gating, reactivation, and reevaluation of economic value in orbitofrontal cortex. Nature Communications 15(1), 6163 (2024)

[40] Ferro, D., Azab, H., Hayden, B.Y., Moreno-Bote, R.: Accumulation of virtual tokens towards a jackpot reward enhances performance and value encoding in dorsal anterior cingulate cortex. Nature Communications 17(1), 7554 (2026)

[41] Oesch, L.T., Thomas, M.C., Sandberg, D., Couto, J., Churchland, A.K.: Anterior cingulate neurons combine outcome monitoring of past decisions with ongoing movement signals. Nature Communications (2026)

[42] Ferro, D., Kempen, J., Boyd, M., Panzeri, S., Thiele, A.: Directed information exchange between cortical layers in macaque v1 and v4 and its modulation by selective attention. Proceedings of the National Academy of Sciences 118(12), 2022097118 (2021)

[43] Bastos, A.M., Usrey, W.M., Adams, R.A., Mangun, G.R., Fries, P., Friston, K.J.: Canonical microcircuits for predictive coding. Neuron 76(4), 695–711 (2012)

[44] Brainard, D.H., Vision, S.: The psychophysics toolbox. Spatial vision 10(4), 433–436 (1997)

[45] Eastman, K.M., Huk, A.C.: Pldaps: a hardware architecture and software toolbox for neurophysiology requiring complex visual stimuli and online behavioral control. Frontiers in neuroinformatics 6, 1 (2012)

[46] Katz, L.N., Yates, J.L., Pillow, J.W., Huk, A.C.: Dissociated functional significance of decision-related activity in the primate dorsal stream. Nature 535(7611), 285–288 (2016)

[47] Bair, W., Movshon, J.A.: Adaptive temporal integration of motion in direction-selective neurons in macaque visual cortex. Journal of Neuroscience 24(33), 7305–7323 (2004)

[48] Van Essen, D.C., Newsome, W.T., Maunsell, J.H.: The visual field representation in striate cortex of the macaque monkey: asymmetries, anisotropies, and individual variability. Vision research 24(5), 429–448 (1984)

[49] Cranmer, K., Brehmer, J., Louppe, G.: The frontier of simulation-based inference. Proceedings of the National Academy of Sciences 117(48), 30055–30062 (2020)

[50] Boelts, J., Lueckmann, J.-M., Gao, R., Macke, J.H.: Flexible and efficient simulation-based inference for models of decision-making. Elife 11, 77220 (2022)

[51] Acerbi, L., Ma, W.J.: Practical bayesian optimization for model fitting with bayesian adaptive direct search. Advances in neural information processing systems 30 (2017)

[52] Singh, G.S., Acerbi, L.: PyBADS: Fast and robust black-box optimization in python. Journal of Open Source Software 9(94), 5694 (2024) 10.21105/joss.05694

[53] Molano-Mazon, M., Barbosa, J., Pastor-Ciurana, J., Fradera, M., Zhang, R.-Y., Forest, J., Pozo Lerida, J., Ji-An, L., Cueva, C.J., Rocha, J., al.: NeuroGym: An open resource for developing and sharing neuroscience tasks. PsyArXiv (2022). 10.31234/osf.io/aqc9n

[54] Brockman, G., Cheung, V., Pettersson, L., Schneider, J., Schulman, J., Tang, J., Zaremba, W.: Openai gym, june 2016. arXiv preprint arXiv:1606.01540 (2016) 10.48550/arXiv.1606.01540

[55] Paszke, A., Gross, S., Massa, F., Lerer, A., Bradbury, J., Chanan, G., Killeen, T., Lin, Z., Gimelshein, N., Antiga, L., Desmaison, A., Kopf, A., Yang, E., DeVito, Z., Raison, M., Tejani, A., Chilamkurthy, S., Steiner, B., Fang, L., Bai, J., Chintala, S.: PyTorch: An Imperative Style, High-Performance Deep Learning Library. Curran Associates, Inc. (2019)

